# *Wt1* splice isoforms bias lineage programmes during formative pluripotency

**DOI:** 10.64898/2026.04.06.713568

**Authors:** Luis Miguel Cerron-Alvan, Tatiana Firfa, Mattia Pitasi, Anne Löbker, Michelle Huth, Michael Bindl, Leonie Drakos, Triin Laos, Martin Leeb

## Abstract

How transcriptional asymmetry first arises in the epiblast prior to gastrulation remains a central question in mammalian development. During the transition from naïve to formative pluripotency, epiblast cells acquire responsiveness to lineage-inducing cues while already expressing lineage-associated transcriptional programmes. The molecular basis of these asymmetries and whether they reflect stochastic variation or regulated lineage-associated bias remain unclear. Here, using a targeted CRISPRa screen, we identify *Wt1* (Wilms tumor 1) as a previously unrecognized regulator of lineage-associated transcriptional programmes during formative pluripotency. *Wt1* is transiently induced during formative pluripotency *in vitro* and in the pre-gastrulation epiblast *in vivo*, with peak expression coinciding with the emergence of lineage-associated transcriptional biases. Precocious *Wt1* induction overrides the naïve transcriptional network and advances cells toward a post-implantation epiblast identity, whereas *Wt1*-deficient cells remain capable of naïve exit but show impaired acquisition of formative and lineage-associated transcriptional programmes. Genome-wide binding analyses during formative state acquisition show that WT1 engages active regulatory elements of the emerging post-implantation gene regulatory network together with core formative transcription factors, including FOXO1, OTX2 and OCT4. Direct comparison of all four major *Wt1* splice isoforms identifies KTS splice status as the major determinant of divergent anterior/neuroectodermal and posterior/mesodermal transcriptional programmes *in vitro*. In the E5.5 epiblast, *Wt1* expression and splice composition align with lineage-biased transcriptional states, linking isoform usage to anterior transcriptional biases *in vivo*. Isoform-dependent gene expression modules are conserved in human pluripotent cells, indicating that this regulatory logic is preserved across species. Together, our findings indicate that lineage-associated transcriptional programmes diverge during formative pluripotency prior to gastrulation and identify alternative splicing as a mechanism that tunes lineage-associated transcriptional bias.

## INTRODUCTION

Pluripotency in the mammalian epiblast unfolds as a continuum, progressing from naïve toward formative and subsequently to primed pluripotency^1–3^. A key unresolved question is whether transcriptional states that bias later lineage outcomes are already encoded within the formative epiblast prior to overt lineage commitment^1,2^.

*In vitro* systems that stabilize the naïve state using two specific inhibitors (2i) and permit synchronized exit upon inhibitor withdrawal have enabled detailed mapping of this progression^4,5^. However, despite these advances, the regulators that actively drive formative state acquisition and shape lineage-associated transcriptional programmes remain incompletely defined.

Most genetic dissection of pluripotency transitions has relied on loss-of-function approaches, which establish necessity but do not test sufficiency. Consequently, the instructive capacity of factors implicated in naïve exit remains less well defined. While several factors have been implicated in naïve exit^6–9^, comparatively few have been shown to function as instructive regulators capable of actively directing cells toward a post-implantation fate. For example, OTX2 has been shown to redirect enhancer landscapes during the transition from naïve to formative pluripotency, yet its instructive capacity under naïve-stabilizing conditions appears limited^10^, suggesting that additional fate-progression determinants remain to be identified.

CRISPR-based transcriptional activation (CRISPRa) provides a strategy to interrogate sufficiency by activating endogenous loci within their native chromatin context^11–14^. Applied under naïve-maintaining conditions, such an approach offers a stringent framework to identify factors capable of overriding the naïve gene regulatory network and advancing post-naïve identity.

To focus this search, we leveraged a defined set of formative-induced genes (FIGs), previously identified as transcripts induced during acquisition of formative identity and inversely correlated with the core naïve network across a large number of perturbations^9^. FIGs represent candidate components of the formative programme, yet their instructive capacity has not been systematically tested.

Here, using a targeted CRISPRa screen in naïve mouse embryonic stem cells, we identify *Wt1* as an early regulator acting during the peri-implantation transition. *Wt1* is transiently induced as cells acquire formative pluripotency and promotes progression toward post-implantation epiblast identity, coinciding with the emergence of lineage-associated transcriptional asymmetry. Precocious *Wt1* expression is sufficient to accelerate progression toward post-implantation epiblast identity, whereas *Wt1* loss impairs normal formative transcriptional progression without preventing naïve exit. *Wt1* is best known for its roles in mesodermal lineage specification and organogenesis, particularly in renal and gonadal development, with no established function in pre-gastrulation pluripotency^15–23^. Notably, alternative *Wt1* splicing generates isoforms that differ by inclusion of a three-amino-acid KTS motif and are associated with distinct lineage-biased transcriptional programmes. *In vitro*, isoforms lacking the KTS motif preferentially engage neuroectodermal-linked programmes, whereas isoforms containing the motif preferentially engage mesodermal-linked programmes. Consistent with this, in the E5.5 epiblast, higher *Wt1* expression is detected in proximity to the emerging anterior visceral endoderm (AVE), while increased representation of transcripts lacking the KTS motif is associated with stronger anterior-linked transcriptional features. Together, our findings identify *Wt1* as an isoform-tuned regulator that shapes lineage-associated transcriptional bias during formative pluripotency.

## RESULTS

### Precocious *Wt1* expression accelerates progression toward post-naïve epiblast identity

To identify regulators that may shape early lineage-associated transcriptional bias during formative pluripotency, we previously identified 218 FIGs which are tightly anticorrelated with the naïve network across more than 100 genetic or chemical perturbations and are upregulated during formative state acquisition^9^ (Supplementary Fig. S1a). We reasoned that this gene set may contain instructive regulators of formative progression. To test this, we performed a targeted CRISPRa screen, using a custom sgRNA library, to activate FIGs in naïve mouse embryonic stem cells (mESCs) maintained in N2B27-2i (N2i) conditions (Fig. 1a). We used Rex1::GFPd2 reporter cells (Rex1-GFP), in which destabilized GFP faithfully tracks transcriptional shutdown of Rex1 during the exit from naïve pluripotency^5,24^, and engineered them to express a doxycycline (dox)-inducible dCas9-VPR transgene, enabling controlled activation of endogenous FIG-loci (Supplementary Fig. S1b).

**Figure 1.**
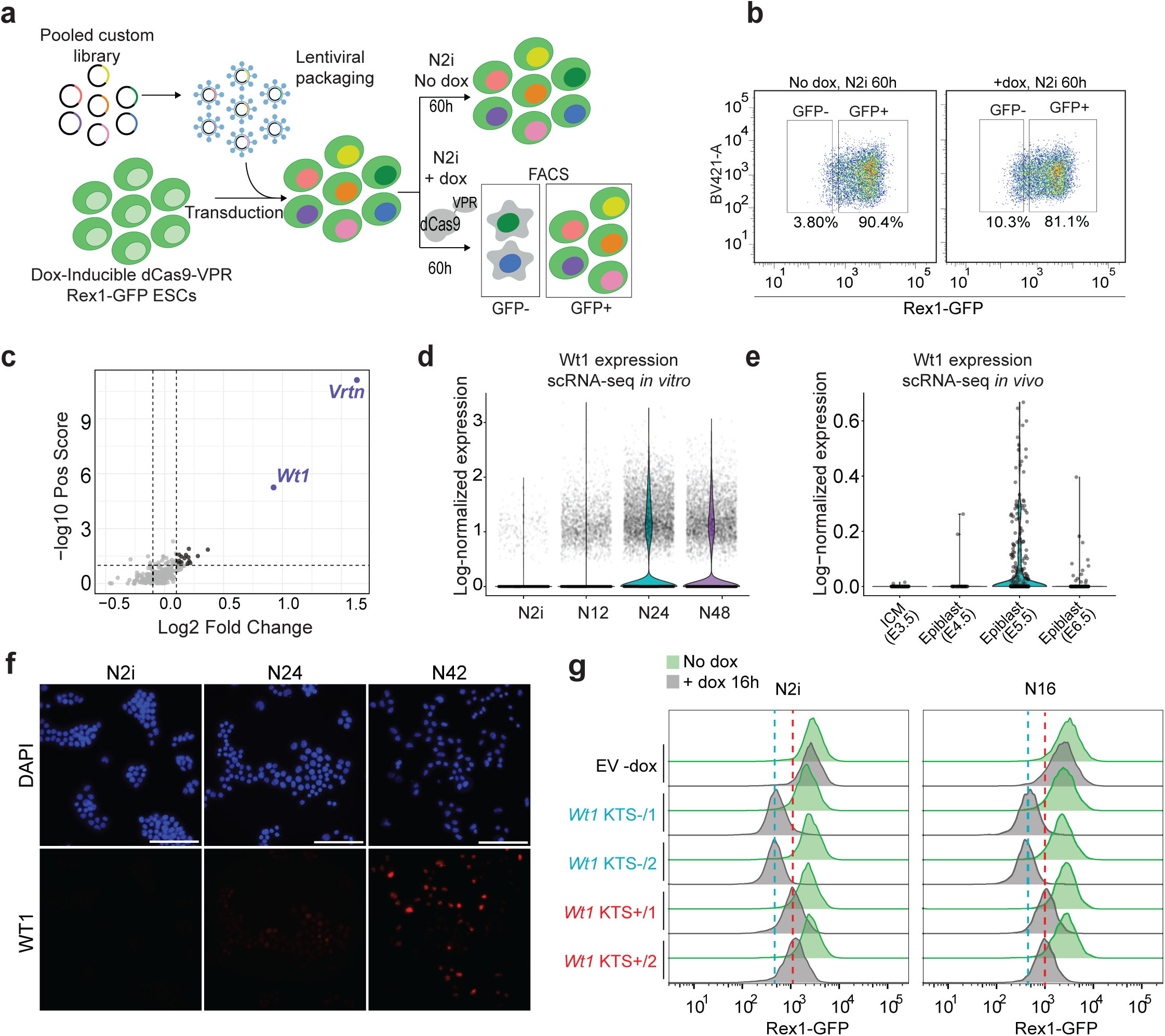
Precocious *Wt1* expression accelerates progression toward post-naïve epiblast identity a. Design of the CRISPRa screen targeting formative induced genes (FIGs). b. Representative flow cytometry showing Rex1-GFP levels in +dox (CRISPRa on) and -dox (off) conditions in 2i. One representative of n = 3 independent experiments is shown. c. MAGeCK gene-level analysis of sorted fractions. Volcano plot showing screen hits enriched in the GFP negative population. d. Single-cell RNA-seq during the in vitro naïve-to-formative transition showing progressive upregulation of *Wt1* expression upon acquisition of the formative state. Nx indicates hours after 2i withdrawal in N2B27. e. *Wt1* expression levels in an integrated in vivo mouse single-cell RNA-seq atlas spanning inner cell mass E3.5 to epiblast stages E4.5, E5.5 and E6.5 (n = 46) from published dataset^26^. f. Immunofluorescence for WT1 in mESCs differentiated from 2i to N24 and N42. Scale bar, 100 μm. g. Inducible precocious expression of the four *Wt1* splice isoforms, *Wt1* KTS-/1, *Wt1* KTS-/2, *Wt1* KTS+/1 and *Wt1* KTS+/2, in clonal lines. Representative Rex1-GFP flow cytometry profiles in 2i and at N16 following 16 h of doxycycline per isoform induction are shown.

After library transduction and 60 h of doxycycline induction in N2i, cells were sorted based on GFP intensity into GFP-positive and GFP-negative fractions for sgRNA quantification. CRISPRa induction increased the proportion of GFP-negative cells compared to non-induced controls, indicating that activation of specific FIGs promotes loss of naïve identity (Fig. 1b). MAGeCK analysis identified multiple genes significantly enriched in the GFP-negative population (Fig. 1c; Supplementary Table 1). Among the top hits, *Wt1* was particularly intriguing, as it is well known for roles in organogenesis and tumorigenesis^20–23^ but has not been implicated in pre-gastrulation pluripotency.

*Wt1* is not detectably expressed in 2i. Upon entering the formative state, however, *Wt1* transcription increases sharply, peaking between N36 and N48 before declining thereafter (Nx indicates x hours after 2i withdrawal in N2B27 medium; Supplementary Fig. S1c)^9,25^. Single-cell RNA-seq confirmed *Wt1* upregulation in a substantial number of cells during monolayer differentiation (Fig. 1d). Consistent with this *in vitro* pattern, analysis of peri-implantation embryo datasets revealed progressive *Wt1* induction from the inner cell mass at E3.5 to highest expression in the post-implantation epiblast at E5.5, followed by downregulation at E6.5 (Fig. 1e)^26^. In line with transcriptional changes observed for *Wt1*, WT1 protein expression peaked at N42, detected by western blot (Supplementary Fig. S1d) and clear but heterogeneous nuclear accumulation in approximately 50% of cells observed by immunofluorescence during the naïve to formative transition (Fig. 1f). *Wt1* encodes multiple splice isoforms that differ primarily in exon 5 inclusion and in alternative splicing of a three–amino acid KTS motif between zinc fingers 3 and 4. The -KTS isoform has been reported to display stronger sequence-specific DNA binding and transcriptional activation, whereas the +KTS isoform shows reduced affinity to DNA^27–31^. To facilitate comparison of the four major *Wt1* splice configurations, we refer to the -KTS/-Ex5, -KTS/+Ex5, +KTS/+Ex5 and +KTS/-Ex5 isoforms as *Wt1* KTS-/1, *Wt1* KTS-/2, *Wt1* KTS+/1 and *Wt1* KTS+/2, respectively. All four splice configurations are expressed during the transition to formative pluripotency, with an overall +KTS:-KTS ratio of approximately 60:40 at peak expression (Supplementary Fig. S1e).

To assess whether precocious *Wt1* expression is indeed sufficient to accelerate progression toward post-naïve epiblast identity, we generated Rex1-GFP clones expressing the four major *Wt1* splice isoforms, *Wt1* KTS-/1, *Wt1* KTS-/2, *Wt1* KTS+/1 and *Wt1* KTS+/2, using a dox-inducible system (Supplementary Fig. S1f). Sixteen hours of forced expression (FE) in 2i conditions resulted in a marked reduction of Rex1-GFP fluorescence for all four isoforms, demonstrating that *Wt1* expression extinguishes naïve identity even under self-renewal-supporting conditions. *Wt1* induction in differentiation-permissive N2B27 medium further accelerated Rex1 downregulation (Fig. 1g), without altering cell morphology or viability (Supplementary Fig. S1g). To assess the effect of prolonged forced *Wt1* expression, we extended doxycycline induction under 2i conditions to 48 h. Under these conditions, both *Wt1* KTS-isoforms maintained an almost complete loss of Rex1-GFP and adopted a more dispersed, elongated morphology. *Wt1* KTS+ expression also reduced Rex1-GFP, although less strongly than the KTS-isoforms, and cells formed flatter, irregular colonies with prominent cellular protrusions (Supplementary Fig. S1h, i).

These results indicated that the strongest inductive potential was determined by absence or presence of the KTS-motif, with exon 5 inclusion showing a minor effect. Therefore, we focused subsequent analyses on two representative isoforms *Wt1* KTS-/1 and *Wt1* KTS+/1, hereafter referred to as *Wt1* KTS- and *Wt1* KTS+, respectively, unless otherwise indicated. Together, these observations identify *Wt1* as a regulator of formative pluripotency that accelerates progression toward post-naïve epiblast identity, while raising the possibility that *Wt1* splice isoforms may exert distinct regulatory functions.

### *Wt1* engages promoters and enhancers of developmental regulators in formative pluripotent cells

To determine the potential regulatory targets of WT1 in the formative state, we performed chromatin immunoprecipitation followed by sequencing (ChIP-seq) during formative state acquisition. Consistent with the increase in protein levels, we detected 236 peaks at N24 in vicinity of 208 protein-coding genes, while at N42, 7,921 peaks close to 4,525 protein coding genes were identified. Virtually all N24 peaks were included in the N42 peak-set (Fig. 2a, b).

**Figure 2.**
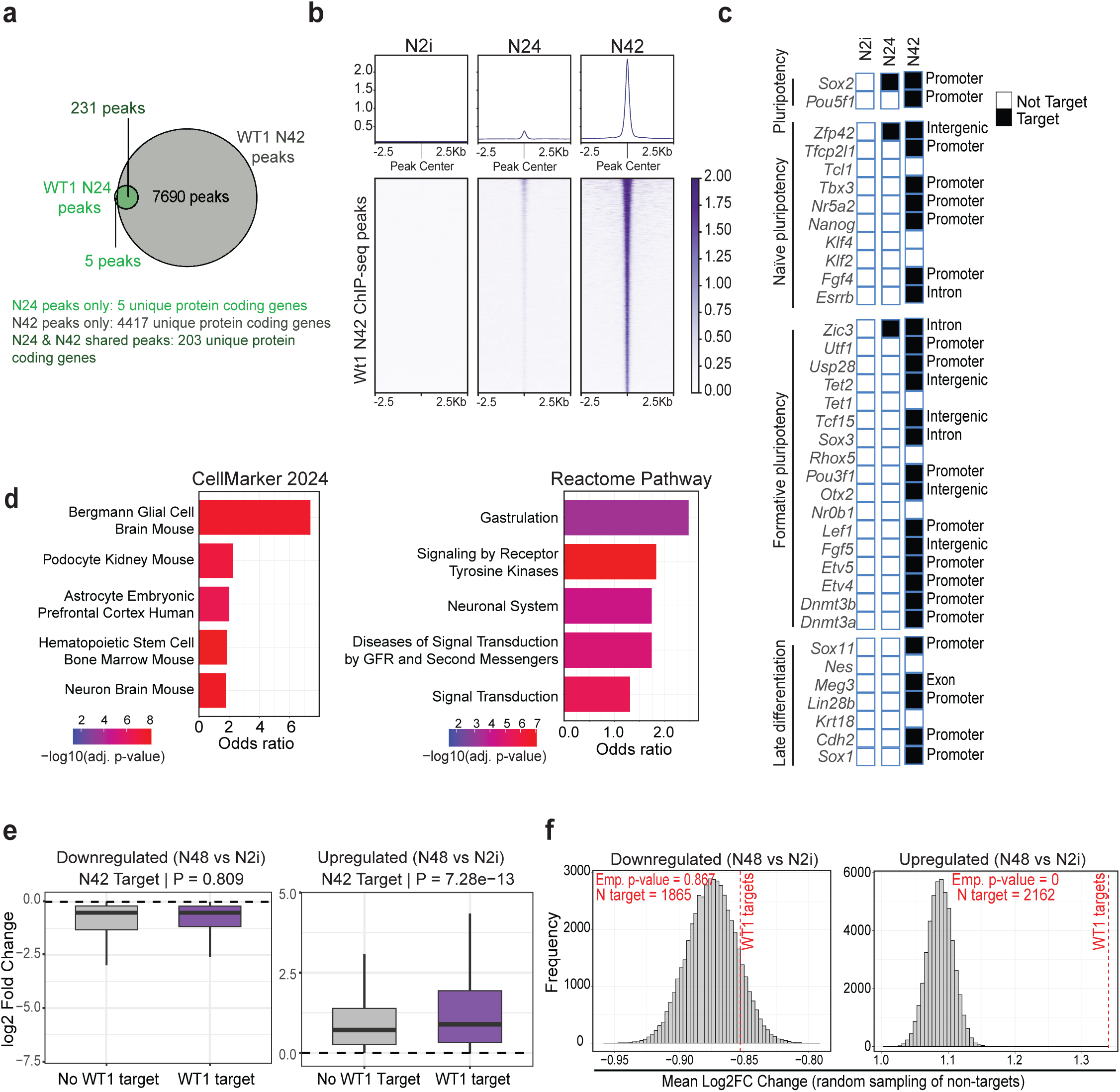
WT1 engages promoters and enhancers of developmental regulators in formative pluripotent cells a. Venn diagram showing overlap of WT1 ChIP-seq peaks at N24 and N42. b. Heatmap of WT1 ChIP-seq signal across peaks at N2i, N24, and N42, illustrating progressive binding upon acquisition of formative pluripotency. c. Representative naïve, formative, and late differentiation marker genes showing WT1 binding dynamics. Genomic annotations were assigned using ChIPseeker. d. Functional enrichment analysis of protein-coding WT1 target genes identified at N42 using Enrichr with Reactome Pathways 2024 and CellMarker gene-set libraries. Stacked bar plots showing the top five enriched terms per library (adjusted p-value ≤ 0.05), ranked by adjusted p-value (ties resolved by overlap count). Bar length indicates odds ratio, and colour scale indicates -log_10_(adjusted p-value). e. Boxplots comparing DESeq2 log_2_ fold changes (N48 vs N2i in wild type) between WT1 target genes and non-target genes, shown separately for upregulated (log2FC > 0) and downregulated (log_2_FC < 0) genes. Group differences were assessed using a two-sided Wilcoxon rank-sum test; p-values are indicated. Sample sizes: downregulated genes, non-target n = 6338, WT1 target n = 1865; upregulated genes, non-target n = 6129, WT1 target n = 2162. f. Permutation analysis of WT1 target gene effects. The observed mean log_2_FC of WT1 targets was compared to null distributions generated from randomly sampled non-target gene sets matched in size (20,000 iterations). Histograms show null distributions, with dashed lines indicating observed means.

At N42, WT1 bound a large set of core peri-implantation cell fate regulators, including naïve (e.g. *Nanog*, *Esrrb*), formative (e.g. *Otx2*, *Pou3f1*), and post-implantation (e.g. *Sox1*, *Chd2*) genes (Fig. 2c; Supplementary Fig. S2a). The majority of peaks was assigned to proximal promoters (27.1%). A substantial fraction of WT1 peaks also mapped to enhancers active during the naïve to formative transition, but not to ESC specific enhancers (19.01% vs 1.83%; Supplementary Fig. S2b).

CellMarker and Reactome enrichment analyses revealed significant neural and glial signatures among WT1-bound genes, alongside gastrulation-related and kidney-associated categories, indicating that WT1 is associated with both ectodermal and mesodermal programmes during formative pluripotency despite its well-established roles in mesodermal lineages (Fig. 2d).

To assess the functional consequence of WT1 chromatin occupancy, we compared expression dynamics of WT1 targets and non-targets during differentiation. While WT1 binding showed little effect on genes downregulated upon formative state acquisition, WT1-bound genes exhibited significantly greater transcriptional induction relative to non-targets (Fig. 2e, f). This is consistent with a predominantly activating role for WT1 during acquisition of post-implantation identity.

Together, these data position WT1 as a component of the emerging post-implantation gene regulatory network, where it engages promoters and enhancers of major developmental regulators and preferentially associates with genes activated during the transition toward post-implantation identity.

### WT1 co-occupies active post-implantation regulatory elements and cooperatively enhances gene activation

To define the regulatory context of WT1 binding, we assessed global co-occupancy with known formative pluripotency factors. Detailed analysis of chromatin-binding data revealed a prominent overlap of WT1 peaks with OCT4, OTX2, FOXO1 and enhancer-associated chromatin marks in formative pluripotent cells (Fig. 3a, b and Supplementary Fig. S3a–c). Notably, WT1 showed markedly stronger overlap with OCT4 peaks in epiblast-like cells (EpiLCs) than with ESC-bound OCT4, indicating preferential association with the post-implantation OCT4 regulon rather than the naïve network (Fig. 3b and Supplementary Fig. S3c). Similar enrichment was observed for OTX2 and FOXO1, placing WT1 within the core post-implantation regulatory circuitry.

**Figure 3.**
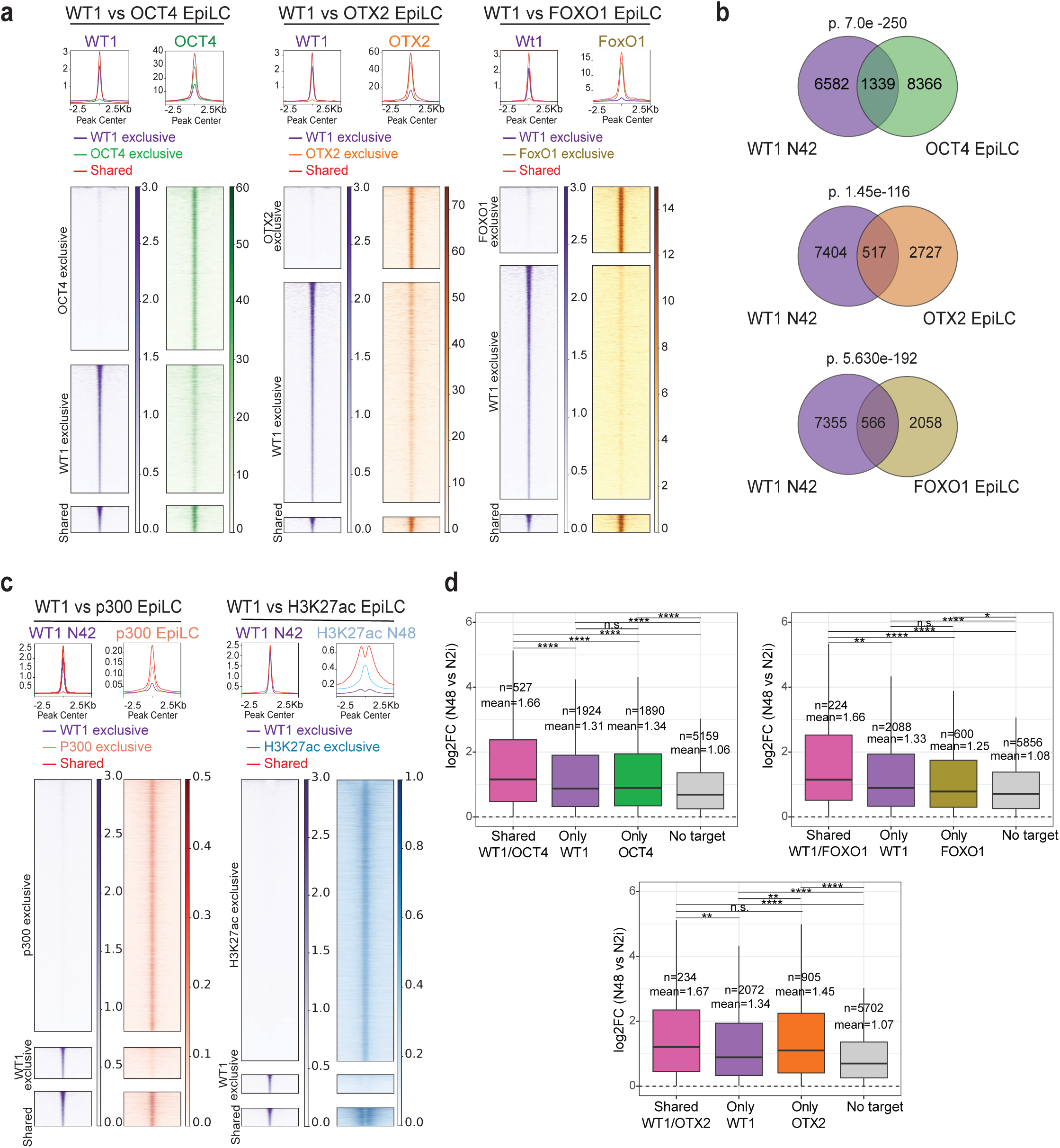
WT1 co-occupies active post-implantation regulatory elements and cooperatively enhances gene activation. a. Heatmaps showing pairwise overlap between WT1 ChIP-seq peaks at N42 and OCT4, OTX2, and FOXO1 ChIP-seq datasets from EpiLCs, with signal plotted ±2.5 kb around peak centres. b. Venn diagrams showing overlap between WT1 peaks at N42 and OCT4, OTX2, and FOXO1 ChIP-seq peaks. Overlap significance was assessed using a hypergeometric test (background n = 155,266 enhancer and open chromatin regions, as in Santini et al.^60^. c. Heatmaps showing overlap between WT1 peaks at N42 and p300 and H3K27ac datasets from formative cells, indicating association with active regulatory elements. Signal is plotted ±2.5 kb around peak centres. d. Boxplots showing expression (log_2_FC, N48 vs N2i) of genes associated with WT1–TF shared peaks, WT1-exclusive peaks, TF-exclusive peaks, and non-target genes. Only genes upregulated during wild-type differentiation were considered. Statistical differences were assessed using pairwise two-sided Wilcoxon rank-sum tests; significance is indicated in the plots (n.s., not significant; *P < 0.05; **P < 0.01; ***P < 0.001; ****P < 0.0001).

Stratification of binding sites into WT1-exclusive, pluripotency TF-exclusive and shared categories revealed that co-occupied regions comprised a significant fraction of WT1 peaks (Fig. 3a,b). Shared sites exhibited higher average ChIP signal-intensities for both WT1 and the partner TF compared to uniquely bound regions, consistent with assembly of high-occupancy regulatory elements at co-bound loci.

WT1-bound regions further extensively overlapped with p300 and H3K27ac peaks in formative cells (Fig. 3c, Supplementary Fig. S3c). Shared WT1-p300 sites exhibited elevated p300 signal compared to p300-only regions, and shared WT1-H3K27ac peaks showed strong H3K27ac enrichment with a central signal-dip characteristic of nucleosome displacement at active enhancers^32^.

We next asked whether co-occupancy translates into functional differences in transcriptional output. Overall, genes linked to shared peaks showed significantly stronger induction during wild-type differentiation, compared to genes associated with WT1-exclusive or TF-exclusive peaks (Fig. 3d). This effect was most pronounced for WT1-OCT4 and WT1-FOXO1 shared loci, with a weaker effect for WT1-OTX2. In contrast, single-bound genes exhibited only modest increases relative to non-targets. These findings indicate that WT1 cooperates with formative transcription factors at shared regulatory elements to enhance gene activation in formative post-implantation epiblast-like cells.

Together, these analyses indicate that WT1 is integrated into active post-implantation regulatory elements co-occupied by OCT4, OTX2 and FOXO1. Co-bound regions exhibit enhanced chromatin activity and are associated with stronger transcriptional induction during the formative transition compared to singly bound loci. These findings position WT1 as a component of the post-implantation gene regulatory network (GRN) that cooperates with formative transcription factors to enhance activation of lineage-associated transcriptional programmes.

### Forced early expression of *Wt1* KTS- and *Wt1* KTS+ accelerates acquisition of post-implantation identity

To define the transcriptional consequences and inductive capacity of precocious *Wt1* expression, we profiled the representative *Wt1* KTS- and *Wt1* KTS+ forced-expression (FE) clones introduced above, which express *Wt1* at levels approximating or below the endogenous peak during formative differentiation (Supplementary Fig. S4a). To further assess expression-level effects, we additionally generated matched higher-expressing lines for both isoforms, designated high expression (HE; Supplementary Fig. S4b). As a reference for an established regulator of the naïve-to-formative transition, we included an inducible *Otx2* high expression line. *Otx2* transcript levels and OTX2 protein abundance peak early during the formative transition around N12 to N16, preceding peak *Wt1* expression (Supplementary Fig. S4c, d). OTX2 promotes formative enhancer engagement by redirecting OCT4 binding during naïve exit^10,24,33^, providing a benchmark for the inductive activity of *Wt1*. In contrast to the pronounced effect of *Wt1* expression on Rex1-GFP levels, even *Otx2* high expression (*Otx2* HE, Supplementary Fig. S4e) failed to reduce Rex1-GFP in 2i and only modestly accelerated Rex1 downregulation under differentiation-permissive conditions (Supplementary Fig. S4f). This indicates that *Wt1* exerts unusually strong instructive effects on post-naïve progression compared to established transition regulators such as *Otx2*.

Transcriptomic profiling of *Wt1* FE cells in 2i and 16 h after 2i withdrawal revealed a pronounced reduction of naïve pluripotency markers accompanied by induction of late formative and committed gene signatures^25^ (Fig. 4a; Supplementary Table 2). These effects were further exacerbated in *Wt1* HE cell lines. By comparison, *Otx2* induction produced only limited remodelling of naïve and early post-implantation regulatory networks under naïve-stabilizing conditions. Under differentiation-permissive conditions, samples with forced *Wt1* expression showed depletion of formative-stage signatures, consistent with accelerated transit through this transient programme toward later post-implantation states. At this resolution, the overall developmental shifts induced by *Wt1* KTS- and *Wt1* KTS+ were largely comparable.

**Figure 4.**
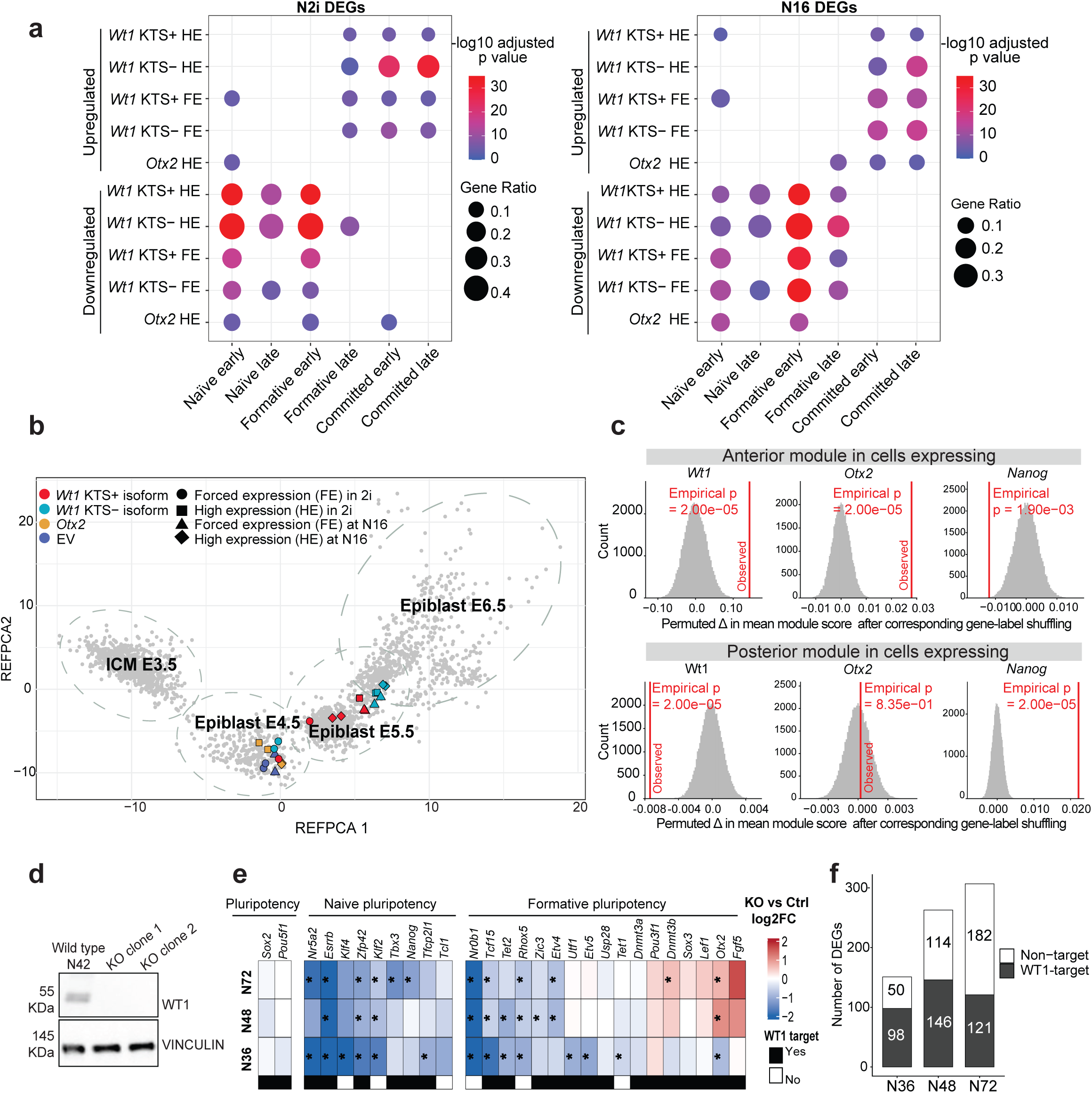
Forced early expression of *Wt1* leads to expedited acquisition of post-implantation identity a. Dot plot showing enrichment of differentially expressed genes (DEGs; padj ≤ 0.05, |log_2_FC| ≥ 0.5) across developmental stage signatures from Carbognin et al.^25^, including naïve (2i), formative (N12–N36), and committed (N48–N72) stages. WT1 conditions comprise forced expression (FE) and high expression (HE). Only significantly enriched categories (adjusted p-value ≤ 0.05) are shown. Multiple testing correction: Benjamini–Hochberg; background: all protein-coding genes. b. Projection of ComBat-corrected bulk RNA-seq profiles onto an integrated single-cell peri-implantation atlas spanning E3.5–E6.5 epiblast development, showing a shift of *Wt1* forced-expression samples toward post-implantation epiblast states. c. Permutation analysis of anterior and posterior module scores in *Wt1*-O*tx2*- and N*anog*-expressing cell populations at N48. Cells were classified as high or low based on log-normalized expression (>0.5, high; ≤0.5, low). For each comparison, group labels were randomly shuffled (50,000 iterations) to generate null distributions of the difference in mean module score. Red lines indicate observed values; empirical p-values are shown. d. Western blot showing WT1 protein levels in wild-type and *Wt1* knockout (KO) clonal lines at N42. VINCULIN serves as a loading control. e. Heatmap showing transcriptional changes in *Wt1* knockout cells during monolayer differentiation. Log_2_ fold changes for selected naïve, formative and lineage-associated differentially expressed genes (DEGs; adjusted p ≤ 0.05, |log_2_FC| ≥ 0.5) are shown for the combined *Wt1* KO clones relative to wild-type controls at N36, N48 and N72. Asterisks indicate statistically significant differential expression. f. Number of differentially expressed genes in *Wt1* knockout cells at N36, N48 and N72, separated into WT1 target and non-target genes. DEGs were defined using adjusted p-value ≤ 0.05 and |log2FC| ≥ 0.5. WT1 targets were defined by WT1 ChIP-seq binding at N42.

To assess how *Wt1* expression alters developmental progression, we integrated FE and HE transcriptomes with *in vitro* differentiation time courses^9,25^ (Supplementary Fig. S4g) and a newly generated integrated single-cell atlas spanning ICM to E6.5 epiblast (Fig. 4b), based on published scRNA-seq datasets^18,26,34–37^. Both *Wt1* isoforms shifted samples along peri-implantation trajectories in an expression-level-dependent manner, with stronger displacement under differentiation-permissive conditions and upon *Wt1* KTS-forced expression. In contrast, *Otx2* high expression did not produce a comparable cell-identity shift under matched conditions.

While trajectory mapping defined developmental identity, it did not resolve GRN-activity states induced by precocious *Wt1* isoform expression. GO analysis of *Wt1*-induced genes revealed an enrichment for neural and morphogenetic processes, with kidney-associated terms more prominently enriched in *Wt1* KTS+ samples (Supplementary Fig. S4h), suggesting emerging isoform-dependent differences within a shared post-implantation framework.

*Wt1*-induced genes significantly overlapped with WT1 ChIP-targets identified at N42 (Supplementary Fig. S4i), indicating that a substantial fraction of the transcriptional response upon forced *Wt1* expression reflects direct WT1 regulatory activity. In the E5.5 embryo, anterior and posterior marker transcripts first become detectable, preceding overt spatial patterning and lineage segregation^26^. This suggests that lineage-associated transcriptional biases may already emerge prior to gastrulation. As N48 in our *in vitro* differentiation system approximates the formative epiblast and corresponds to the E5.5 stage, we analysed anterior and posterior transcriptional modules in wild-type single-cell RNA-seq data obtained at N48. At this stage, *Wt1*, *Otx2* and *Nanog* are heterogeneously expressed across the single-cell population (Supplementary Fig. S4j). We then asked whether cells expressing high *Wt1* levels (*Wt1*-high) preferentially express anterior or posterior gene modules compared to cells expressing low or undetectable *Wt1* levels (*Wt1*-low). As controls we investigated *Otx2*-high and low and *Nanog*-high and low expressing cell populations. *Otx2* is a well-studied anterior marker^38^ and *Nanog* has been shown to be repurposed to shut down pluripotency in the post-implantation epiblast^39^ (Fig. 4c). As expected, *Otx2*-high cells showed a strong enrichment of anterior marker gene expression, whereas *Nanog*-high cells exhibited a significantly stronger expression of posterior marker genes and a reduction of anterior gene expression. Strikingly, *Wt1*-high cells exhibited increased anterior module scores and significantly reduced posterior scores relative to *Wt1*-low cells, despite the well-established later roles of *Wt1* in mesoderm-derived contexts^40^.

To determine whether transcriptional changes during formative state acquisition depend on endogenous *Wt1*, we first analysed two independent *Wt1* knockout clones during monolayer differentiation. Loss of WT1 protein was confirmed at N42 by western blotting (Fig. 4d). Transcriptomic profiling at N36, N48 and N72 showed that *Wt1*-deficient cells remained capable of progressing beyond the naïve state but displayed impaired transcriptional progression during the naïve-to-formative transition. Developmental, formative and lineage-associated genes showed frequently reduced expression in the knockout, with WT1-bound targets prevalent among the affected genes (Fig. 4e,f). Notably, *Otx2* showed an opposite early response. It increased following *Wt1* loss and decreased upon precocious *Wt1* expression (Fig. 4e, Supplementary Table 2). Consistently, genes downregulated following *Wt1* loss were enriched for general developmental processes across the analysed time points (Supplementary Fig. S4k).

Collectively, these data identify *Wt1* as a regulator that advances cells along an authentic post-naïve developmental trajectory during formative pluripotency, while exhibiting a pronounced and unexpected bias toward anterior transcriptional programmes.

### *Wt1* splice isoforms bias lineage programmes during early differentiation

To investigate isoform-specific *Wt1* functionality in an unbiased differentiation model, we utilized an embryoid body (EB) system. To this end, mESCs carrying inducible *Wt1* KTS- or *Wt1* KTS+ transgenes were aggregated into EBs and were grown for 3 days under constant *Wt1*-induction prior to transcriptomic analysis (Supplementary Fig. S5a, b; Supplementary Table 2).

Both isoforms induced substantial transcriptional changes in EBs at day 3 of differentiation, with a significant shared component (Fig. 5a). However, isoform-exclusive gene sets revealed distinct programme-enrichment. Genes uniquely upregulated by *Wt1* KTS-were enriched for neuroectodermal lineage categories, whereas *Wt1* KTS+-exclusive genes were enriched for mesodermal categories (Fig. 5b). Genes induced by both isoforms were enriched for general post-implantation progression signatures, indicating a shared post-implantation fate-progression programme alongside a divergent lineage bias. Complementary profiling of EBs inducibly expressing the *Wt1* KTS-/2 and *Wt1* KTS+/2 isoforms showed that these lineage-associated responses were largely reproduced according to KTS status (Supplementary Fig. S5d). Thus, direct comparison of all four isoforms identifies KTS splice status as the major determinant of the divergent lineage-associated transcriptional phenotype in our assays. Exon 5 inclusion is therefore not required for either the KTS--associated neuroectodermal bias or the KTS+-associated mesodermal bias, although more subtle effects of exon 5 inclusion cannot be excluded.

**Figure 5.**
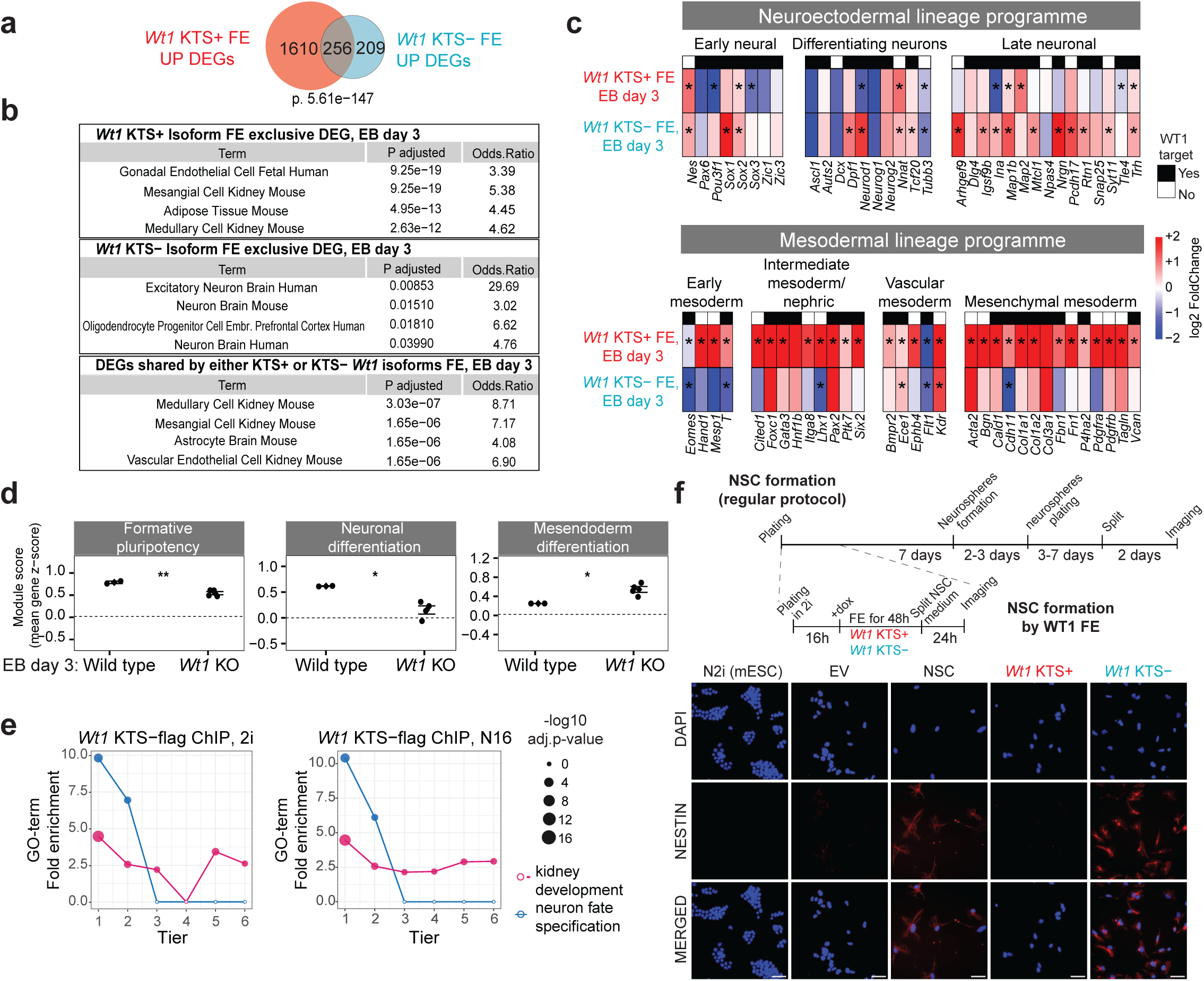
*Wt1* splice isoforms bias lineage programmes during early differentiation a. Transcriptomic comparison *Wt1* KTS+ and *Wt1* KTS-forced expression in embryoid bodies (EBs, day 3). Differentially expressed genes (DEGs; padj < 0.1) identified from two independent clonal lines per isoform are shown as a Venn diagram. FE, forced expression. b. Functional enrichment of isoform-specific transcriptional responses. DEGs exclusive to *Wt1* KTS+, exclusive to *Wt1* KTS-, and shared between isoforms were analysed using CellMarker gene sets (Enrichr); top enriched terms are shown. c. Lineage marker responses to *Wt1* isoform expression. Heatmap showing log_2_ fold changes (EB day 3, vs control) across neuroectodermal and mesodermal lineage programmes. Marker genes were curated and validated against CellMarker databases. Asterisks indicate adjusted p-value < 0.1. *Wt1* target genes (defined by N42 ChIP-seq) are indicated. d. Module-score analysis of lineage-associated transcriptional programmes in *Wt1* knockout embryoid bodies at day 3. Module scores for formative pluripotency, neuronal differentiation and mesendoderm were calculated from TMM-normalized logCPM values after gene-wise z-scoring across samples. Wild-type and *Wt1* KO samples are shown; statistical significance was assessed using Welch’s t-test. e. Functional enrichment across *Wt1* KTS-ChIP-binding tiers. Fold enrichment of GO terms is shown across binding tiers, with terms associated with neuronal and kidney development highlighted. Dot size indicates significance (-log_10_ adjusted p-value). f. Schematic overview of neural stem cell (NSC) differentiation. The upper scheme shows the standard NSC differentiation protocol, while the lower scheme shows the experimental strategy used to assess NSC generation following forced expression of *Wt1* KTS+, *Wt1* KTS-, or empty vector control under differentiation-permissive conditions, followed by culture in NSC medium. Immunofluorescence analysis of the NSC differentiation assay. NESTIN and DAPI staining are shown at endpoint. Scale bar, 50 μm.

Analysis of curated anterior, posterior, neuronal and mesodermal gene sets supported this divergent functionality: *Wt1* KTS-preferentially activated anterior and neuroectoderm-associated programmes, whereas *Wt1* KTS+ predominantly engaged posterior and mesoderm-derived modules while showing relative repression of neural gene sets (Fig. 5c, Supplementary Fig. S5c).

To assess whether these transcriptional programmes are dependent on *Wt1*, we profiled *Wt1* knockout EBs at days 3 and 10 (Supplementary Fig. S5e; Supplementary Table 2). At day 3, *Wt1*-deficient EBs showed reduced formative and neuronal differentiation-associated programme scores together with increased mesendoderm-associated scores (Fig. 5d). Transcriptomic analysis further revealed widespread changes in developmental and lineage-associated gene expression (Supplementary Fig. S5e). By day 10 mesendoderm-associated programme scores were reduced in knockout EBs (Supplementary Fig. S5f). Together, these results reveal stage-dependent consequences of *Wt1* loss, with the early phenotype showing the reciprocal lineage-associated shift predicted from our gain-of-function experiments and the later phenotype consistent with the established functions of *Wt1* in mesoderm-derived developmental programmes.

Direct comparison of forced expression and knockout datasets revealed a pronounced opposing regulation for *Wt1* KTS-. Genes induced upon *Wt1* KTS-expression were frequently reduced in the knockout and vice versa, with the strongest inverse correlation among WT1-bound targets with significant deregulation in both contexts (Supplementary Fig. S5g, h). The strong inverse relationship between *Wt1* KTS-forced expression and global *Wt1* loss, particularly among WT1-bound targets, identifies the KTS--associated programme as a prominent component of endogenous *Wt1* function during formative pluripotency.

Collectively, these analyses indicate that while both KTS classes are sufficient to promote post-implantation progression, the KTS splice configuration encodes qualitatively distinct transcriptional biases. *Wt1* KTS-preferentially engages anterior neural programmes, whereas *Wt1* KTS+ favours posterior mesodermal and kidney-associated gene expression modules. These findings support a model in which *Wt1* splice composition modulates lineage programme engagement during formative pluripotency.

Given the neural-biased transcriptional output observed upon *Wt1* KTS-induction, we next asked whether this isoform preferentially occupies neural programme-associated regulatory loci. To address this, we performed FLAG ChIP-seq following induction of FLAG-tagged *Wt1* KTS-in monolayer culture and compared these profiles with ChIP-seq data obtained for endogenous WT1. High-confidence FLAG-*Wt1* KTS-binding sites were preferentially associated with neural fate specification terms, whereas lower-intensity sites, or sites detected only by the pan-WT1 antibody recognizing endogenous WT1, were increasingly linked to kidney-related categories (Fig. 5e, Supplementary Fig. S5i). This association between *Wt1* KTS-binding intensity and functional annotation was preserved in both 2i and during the transition to formative pluripotency.

To determine whether this binding bias translates into distinct functional lineage outcomes, we performed a directed neural stem cell differentiation assay. We observed that transient *Wt1* KTS-induction markedly accelerated neural stem cell specification, evidenced by rapid acquisition of NSC-like morphology and robust NES (Nestin) expression, which was not evident in *Wt1* KTS+ expressing or in control cells (Fig. 5f, Supplementary Fig. S5j). These findings indicate that the anterior-biased transcriptional programme induced by *Wt1* KTS-is sufficient to accelerate neural lineage specification, consistent with a functionally instructive role in promoting neuroectodermal differentiation.

Collectively, these analyses indicate that while both *Wt1* isoforms promote post-implantation progression, splice configuration encodes qualitatively distinct transcriptional biases. *Wt1* KTS-preferentially engages anterior neural programmes, whereas *Wt1* KTS+ favours posterior mesodermal and kidney-associated modules. These findings support a model in which *Wt1* splice composition modulates lineage programme engagement, with *Wt1* KTS-emerging as the dominant regulator during formative pluripotency.

### *Wt1* isoform programmes show cross-species transcriptional conservation

Endogenous *WT1* is also transiently upregulated during naïve exit in human ESCs (hESC), although expression remains generally low and the increase is less pronounced than the formative-associated *Wt1* wave observed in mouse. In SHEF6 cells, *WT1* expression increased transiently at days 4–6 before declining again by day 8 (Supplementary Fig. S6a)^41^. We therefore asked whether the isoform-dependent regulatory logic identified in mouse is conserved in human pluripotency transitions. To test this, we expressed *Wt1* KTS- or *Wt1* KTS+ in differentiating SHEF6 cells and defined *Wt1*-responsive genes relative to empty-vector controls (Supplementary Fig. S6b).

Partitioning upregulated genes into common, *Wt1* KTS+ specific, and *Wt1* KTS-specific sets revealed a pattern similar to that observed in mouse. The KTS-specific gene set was enriched for neural and glial cell types, whereas the KTS+ specific set preferentially returned kidney-associated and non-neural annotations (Fig. 6a, b). Thus, even in a heterologous human context, the two isoforms exhibited divergent lineage-biased transcriptional outputs.

**Figure 6:**
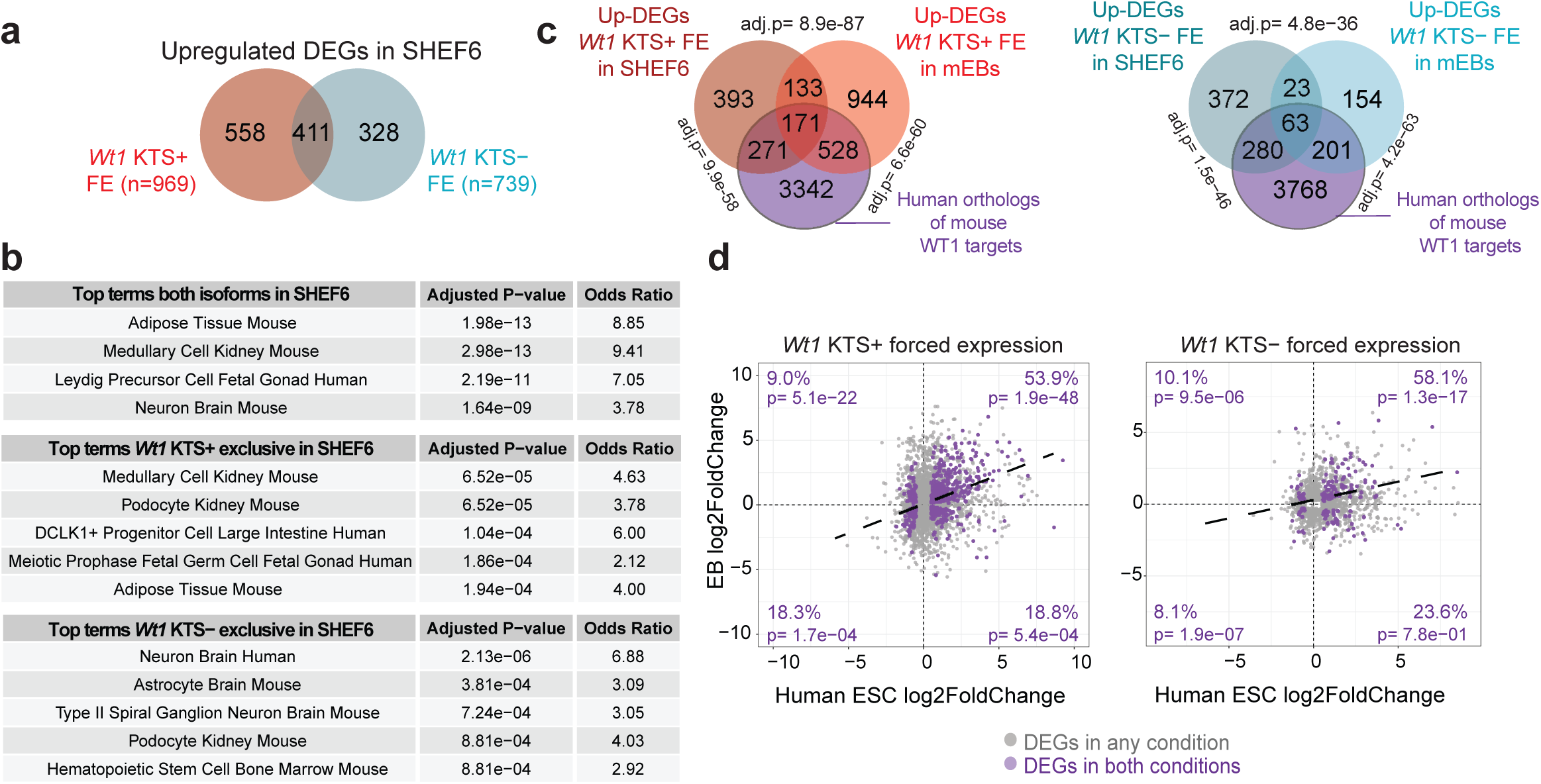
*Wt1* isoform programmes show cross-species transcriptional conservation a. Overlap between *Wt1* KTS– and *Wt1* KTS+ induced upregulated genes in human SHEF6 cells at day 2 of differentiation. Venn diagrams define shared and isoform-specific transcriptional responses (|logFC| ≥ 0.5; adjusted p-value ≤ 0.05). b. Cell-type annotation of isoform-specific and shared gene sets using CellMarker gene sets (Enrichr). Top enriched terms are shown for each category. c. Cross-species comparison of *Wt1*-regulated gene sets. Triple Venn diagrams show overlap between human *Wt1* forced expression DEGs (SHEF6, day 2), mouse embryoid body DEGs (EB day 3), and *Wt1* target gene orthologs, shown separately for *Wt1* KTS+ and *Wt1* KTS-. Mouse genes were mapped to human using strict 1:1 orthology. Enrichment significance was assessed using hypergeometric tests. d. Concordance of transcriptional responses across species. Scatterplots compare gene-wise log fold changes between human (SHEF6) and mouse (EB) *Wt1* isoform forced expression for 1:1 orthologs. Genes significant in both datasets are highlighted. For common DEGs only, deviation from an expected 25:25:25:25 quadrant distribution was tested by χ² goodness-of-fit, yielding χ² p = 4.1e-57 for *Wt1* KTS+ and χ² p = 1.87e-20 for *Wt1* KTS-. Quadrant-specific enrichment or depletion was evaluated using two-sided exact binomial tests against an expected proportion of 0.25. Quadrant labels indicate the corresponding percentages and two-sided exact binomial test p-values. Purple points and the dark purple fitted line indicate the common DEG subset.

Mapping human DEGs to mouse orthologs demonstrated a significant overlap between isoform-matched DEG-sets in differentiating hESCs and mouse embryoid bodies (Fig. 6c). Shared DEG subsets showed a pronounced and highly significant concordant expression change, indicating that common *Wt1*-responsive genes were preferentially upregulated in mouse and human formative pluripotency models (Fig. 6d). This effect was observed for both isoforms, with more than half of shared DEGs showing concordant changes in expression (*Wt1* KTS+, 53.9% up-up; *Wt1* KTS-, 58.1% up-up), suggesting conservation of a core *Wt1*-associated activation-programme across species.

### *Wt1* expression and splice composition align with lineage-biased transcriptional states in the E5.5 epiblast

To determine whether the *Wt1*-responsive regulatory programmes identified *in vitro* are also active *in vivo*, we examined transcriptional states of *Wt1*-high and *Wt1*-low cells in the E5.5 epiblast. We first focused on a set of 595 WT1-bound target genes (*Wt1*-target module) that were significantly upregulated upon forced *Wt1* expression in EBs. Scoring this gene module across single E5.5 epiblast cells revealed a significantly higher *Wt1* target module score in *Wt1*-high cells compared with *Wt1*-low cells, a difference corroborated by permutation testing (Supplementary Fig. S7a). Thus, the *Wt1*-responsive target programme identified *in vitro* is preferentially active in *Wt1*-high cells of the E5.5 epiblast.

We next asked whether *Wt1*-high expressing cells in the E5.5 epiblast exhibit a transcriptional bias toward anterior or posterior programmes. To this end, we compared the expression of anterior and posterior GRN modules between *Wt1*-high and *Wt1*-low expressing cells; as controls, we analyzed *Otx2*-high and *Otx2*-low, and *Nanog*-high and *Nanog*-low expressing cells (Fig. 7a; Supplementary Figs. S7b, c). This analysis revealed the expected anterior bias of *Otx2*-high expressing cells. *In vivo* at E5.5, *Nanog*-high expressing cells exhibited both anterior and posterior transcriptional features, potentially reflecting population-level heterogeneity in the formative epiblast. *Wt1*-high cells showed a significant enrichment of anterior programmes, similar in magnitude to that observed for *Otx2*. Thus, *Wt1*-high cells in the E5.5 epiblast are characterized by stronger anterior-associated transcriptional programme activity prior to gastrulation.

**Figure 7.**
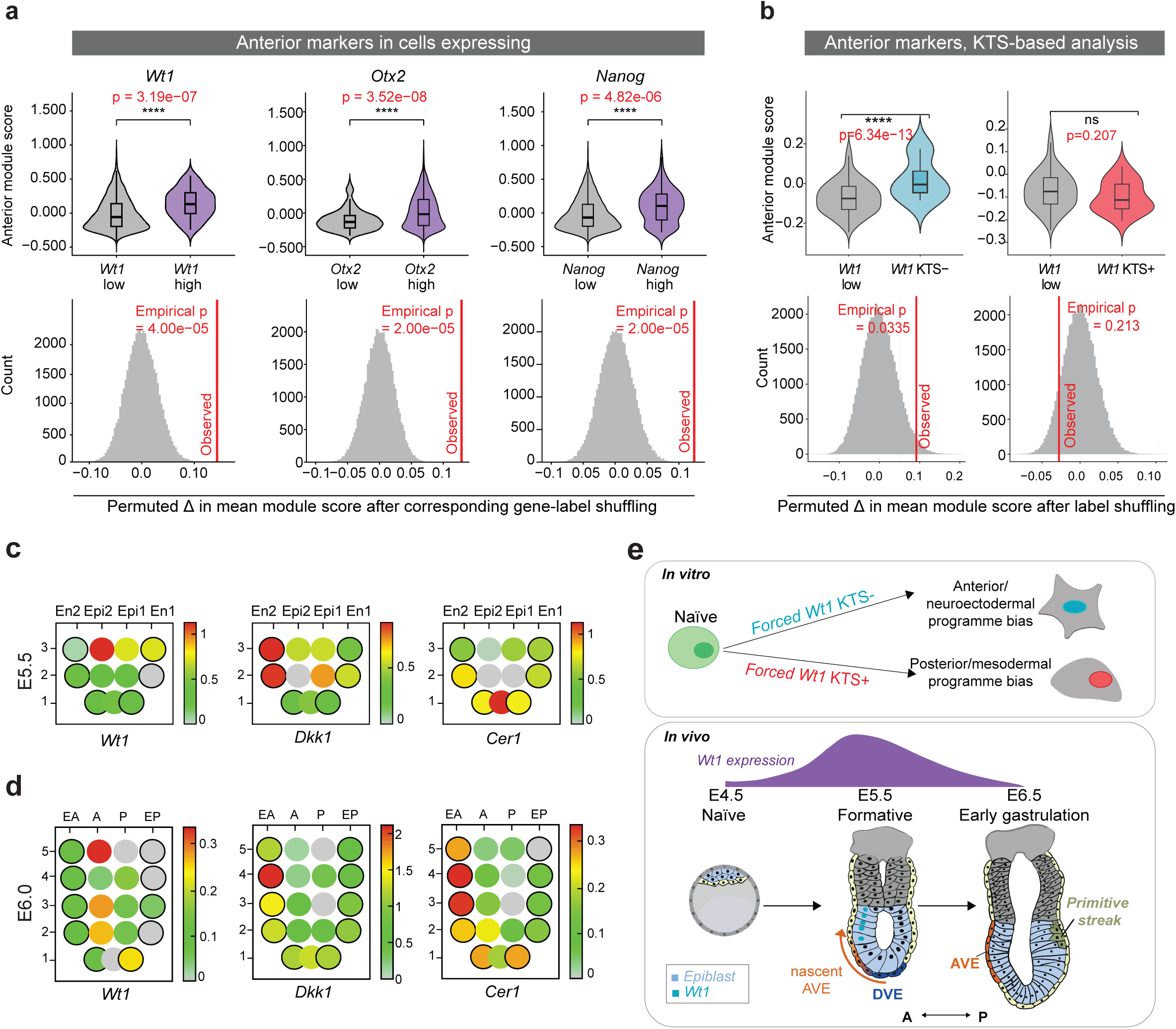
*Wt1* expression and splice composition align with lineage-biased transcriptional states in the E5.5 epiblast a. Violin plots and permutation analyses comparing anterior module scores between *Wt1*-low and *Wt1*-high, *Otx2*-low and *Otx2*-high, and *Nanog*-low and *Nanog*-high E5.5 epiblast cells. For each comparison, group labels were shuffled 50,000 times to generate null distributions of the difference in mean module score (Δ); vertical red lines indicate the observed Δ. b. Violin plots and permutation analyses comparing cells with predominant *Wt1* KTS- or *Wt1* KTS+ with *Wt1*-low cells for anterior module scores using published E5.5 epiblast Smart-seq2 single-cell RNA-seq dataset (266 E5.5 cells)^26^, which enabled splice-isoform-resolved analysis. Group labels were shuffled across cells 50,000 times to generate null distributions, and Δ was recalculated for each permutation; vertical red lines indicate the observed Δ. c. Spatial expression of *Wt1*, *Dkk1* and *Cer1* in E5.5 mouse embryos using the eGastrulation spatial transcriptomic dataset GSE120963. Corn plots retain the spatial organization and regional nomenclature of the reference embryos defined in the original dataset. Each row represents a spatial level along the embryo, with section numbers corresponding to the original GEO-seq sampling scheme. At E5.5, Epi1 and Epi2 denote the two epiblast regions and En1 and En2 the corresponding endoderm regions. Following the original corn-plot convention, solid circles represent ectodermal samples, black-outlined circles represent endodermal samples, and grey hollow circles indicate positions for which no sample was available. Expression values are shown using the colour scale indicated for each gene. d. Similar to c for E6.0. At E6.0, A and P denote anterior and posterior epiblast regions, respectively, while EA and EP denote anterior and posterior endoderm. e. *In vitro*, forced *Wt1* expression promotes exit from naïve pluripotency and progression toward post-implantation transcriptional states. *Wt1* KTS- and *Wt1* KTS+ show divergent lineage-associated activities, biasing anterior/neuroectodermal and posterior/mesodermal transcriptional programmes, respectively. *In vivo*, *Wt1* expression is low in the naïve epiblast at E4.5, increases transiently around E5.5, and declines by E6.5. At E5.5, higher *Wt1* expression is associated with stronger anterior programme activity, while predominant KTS-representation is associated with increased anterior and neural stem-cell programme scores, consistent with the KTS--associated bias observed *in vitro*. DVE, distal visceral endoderm; AVE, anterior visceral endoderm; A, anterior; P, posterior.

Given the isoform-specific lineage biases observed *in vitro*, we next asked whether endogenous variation in *Wt1* splice composition within E5.5 epiblast cells is associated with similar lineage-linked transcriptional tendencies. To this end, we leveraged available full-length SMART-seq2 data and computed module scores for anterior and posterior patterning-related genes and examined their relationship to *Wt1* splice features. Cells were grouped according to the predominant KTS splice configuration among informative junction reads; this classification reflects relative splice representation rather than mutually exclusive expression of KTS- or KTS+ transcripts. Predominant KTS splice representation was associated with clear shifts in the anterior-associated programme (Fig. 7b). Cells with predominant *Wt1* KTS-representation exhibited significantly increased anterior-associated expression while showing reduced posterior module expression. In contrast, cells with predominant *Wt1* KTS+ representation did not show detectable enrichment of lineage-specific transcriptional programmes. Consistent with this, cells with predominant KTS-representation exhibited increased anterior neural stem cell module scores and reduced posterior nephron progenitor module scores (Supplementary Fig. S7d). These findings indicate that variation in *Wt1* splice composition is associated with lineage-associated transcriptional biases within the E5.5 epiblast, consistent with the isoform-specific biases observed *in vitro*.

We next examined the spatial relationship between *Wt1* expression and emerging anterior-posterior patterning using published GEO-seq data from peri-implantation mouse embryos^42^. At E5.5, higher *Wt1* expression was associated with the side of the embryo showing stronger expression of the AVE-associated markers *Dkk1* and *Cer1* (Fig. 7c). By E6.0 overall *Wt1* expression was reduced, with the remaining signal showing a more anteriorly localized distribution (Fig. 7d). As after E5.5, *Wt1* expression was strongly reduced, we restricted KTS analysis to E5.5 epiblast samples containing informative KTS splice-junction fragments. For each informative sample, we determined the fraction of KTS-transcripts and compared it with a relative anterior-posterior transcriptional score, calculated by subtracting posterior from anterior module activity, with higher values indicating stronger relative anterior-associated gene expression. Samples with a higher KTS-fraction tended to display higher anterior-minus-posterior scores, associating increased KTS-representation with a more anterior-biased transcriptional state (Supplementary Fig. S7e). Together, these spatial analyses show that *Wt1* expression and splice composition are associated with the emerging anterior-posterior asymmetry of the embryo at E5.5.

Together, these analyses show that the *Wt1*-responsive regulatory programme defined functionally *in vitro* is preferentially active in *Wt1*-high cells of the E5.5 epiblast, linking stem-cell perturbation data to endogenous transcriptional variation *in vivo*. Within this population, predominant KTS-representation is associated with stronger anterior and neural-linked programme activity, closely paralleling the bias induced by *Wt1* KTS-in functional *in vitro* assays. By contrast, the posterior/mesodermal activity of *Wt1* KTS+ is supported primarily by our *in vitro* differentiation experiments and is not equivalently resolved *in vivo*. Collectively, these findings connect the KTS--associated transcriptional programme identified functionally *in vitro* with emerging anterior transcriptional asymmetry in the E5.5 epiblast, coincident with peak *Wt1* expression and its spatial association with the nascent AVE (Fig. 7e).

## DISCUSSION

A central question in early mammalian development is when and how lineage-associated transcriptional asymmetry first emerges within the epiblast, and whether these asymmetries reflect stochastic variation or regulated lineage-associated programmes. Our findings suggest that regulated lineage-associated transcriptional biases emerge during formative pluripotency, prior to overt lineage segregation and before gastrulation. This indicates that transcriptional symmetry within the epiblast may already be broken at the level of transcriptional programmes at the formative pluripotent state.

We identify *Wt1* as an early regulator acting during formative pluripotency. *Wt1* expression peaks during the late formative stage *in vitro* and is prominently detected in the E5.5 epiblast *in vivo*, preceding overt lineage segregation. *Wt1* expression declines again as gastrulation begins. In this light, the *Wt1* signal detected in early gastrulation datasets likely reflects the trailing edge of this earlier expression wave. Together, these observations suggest that *Wt1* does not simply act earlier than previously appreciated^19^ but instead marks a previously unrecognized wave of regulatory activity during formative pluripotency.

*Wt1* is classically regarded as a mesoderm-associated developmental regulator required for renal and gonadal development^20,43^. Early expression mapping positioned the onset of *Wt1* function during organogenesis, with transcripts detected in nephrogenic mesoderm from E9.0–E9.5^44^. More recent work has defined the onset of *Wt1* activity earlier in development. Single-cell analyses during gastrulation report *Wt1* expression beginning around E6.5 in primitive streak derivatives and mesenchymal populations associated with posterior lineage formation^19^. *Wt1* activity has primarily been interpreted in the context of mesoderm specification during gastrulation.

In contrast, our data indicate that *Wt1* does not behave as a classical mesoderm determinant but instead engages active regulatory elements of the emerging post-implantation gene regulatory network. At the chromatin level, WT1 is integrated into the formative regulatory circuitry through extensive co-occupancy with OCT4, OTX2 and FOXO1, with genes associated with co-bound regulatory elements showing stronger transcriptional induction than singly bound loci. Alternative *Wt1* splice isoforms encode and bias transcriptional programmes toward anterior neuroectodermal or posterior mesodermal modules, suggesting that *Wt1* acts upstream of lineage specification to influence lineage-associated transcriptional programmes within the formative epiblast. These observations support a model in which early epiblast regulators establish lineage-associated transcriptional biases before overt lineage specification.

The timing of *Wt1* expression indicates that *Wt1* does not trigger naïve exit. Instead, its induction coincides with the transition phase during which cells commit to differentiation and acquire post-implantation epiblast features. *Wt1* reinforces the transition toward post-naïve regulatory states. When expressed precociously, *Wt1* rapidly suppresses naïve identity and engages a post-implantation gene regulatory network within 16 hours, even under naïve-stabilizing 2i conditions. *Otx2*, a canonical transition regulator, did not produce comparable effects under matched conditions, highlighting *Wt1* as a particularly potent regulator of early post-implantation developmental progression. The temporal relationship between *Otx2* and *Wt1* further suggests that these regulators act at distinct phases of formative progression. *Otx2* abundance peaks early during the formative transition, preceding the later peak of *Wt1* expression. *Otx2* is oppositely regulated by *Wt1* perturbation: increasing following *Wt1* loss and decreasing upon precocious *Wt1* expression. Together with the stronger capacity of precocious *Wt1* expression to advance post-naïve identity, these observations support a model in which Otx2 contributes to early formative-state establishment, whereas *Wt1* acts subsequently to promote maturation toward later post-implantation transcriptional states. *Wt1*-deficient cells remain capable of leaving the naïve state but show impaired acquisition of formative and lineage-associated transcriptional programmes. The consequences of *Wt1* loss are stage dependent: early differentiating EBs show reduced formative and neuronal programme activity together with increased mesendoderm-associated activity, whereas at later stages mesendoderm-associated programmes are reduced. Thus, the early contribution of *Wt1* to anterior-associated transcriptional programmes appears specific to the formative state, whereas its later activity is consistent with its established functions in mesodermal development. Despite pronounced sufficiency effects and clear opposing transcriptional responses in gain- and loss-of-function settings, lineage programmes can still be initiated in the absence of *Wt1*, indicating that *Wt1* contributes to their regulation but is not uniquely required, likely reflecting parallel or compensatory pathways within the peri-implantation regulatory network, similar to the cooperative and partially redundant genetic control of exit from naïve pluripotency^9,45^.

A distinctive feature of *Wt1* biology is the generation of multiple protein isoforms through alternative splicing. The two principal splice events involve inclusion or exclusion of exon 5 and insertion or omission of a KTS tripeptide between zinc fingers 3 and 4^46^. These splice choices influence WT1 molecular behaviour, including DNA-binding properties and nuclear localisation, and altered KTS+/KTS-ratios cause developmental pathology in humans^47,48^. Consistent with functionally distinct roles of the KTS splice variants, *Wt1* KTS-was recently shown to be essential for ovarian determination in mice^43^. Our data show that isoform composition can encode and tune lineage-associated transcriptional outputs. Our direct comparison of all four major *Wt1* splice isoforms identifies KTS-status as the major determinant of the divergent lineage-associated transcriptional response. *Wt1* KTS-isoforms preferentially engage neuroectodermal and neural programmes, whereas *Wt1* KTS+ isoforms more strongly activate mesodermal, stromal and kidney-associated modules. Exon 5 inclusion is not required for either transcriptional bias, although more subtle effects of exon 5 inclusion cannot be excluded. The KTS--associated neural bias is supported at both regulatory and functional levels, with *Wt1* KTS-binding preferentially associated with neural programme genes and transient *Wt1* KTS-expression accelerating neural stem cell specification.

Notably, both KTS splice classes are expressed during the formative transition, with KTS+ transcripts constituting the larger fraction at peak *Wt1* expression, yet their endogenous functional contributions appear unequal. Whereas forced expression reveals distinct inductive capacities of *Wt1* KTS- and *Wt1* KTS+, the transcriptional consequences of global *Wt1* loss align predominantly with the *Wt1* KTS--associated programme. Thus, KTS--associated activity appears to constitute a major component of endogenous *Wt1* function during formative pluripotency, despite the capacity of *Wt1* KTS+ to induce mesoderm-associated programmes upon forced expression.

Evidence from the embryo supports this regulatory logic. The lineage-associated activity identified by forced expression is also reflected in endogenous *Wt1*-associated states. During *in vitro* formative differentiation, *Wt1*-high cells show increased anterior and reduced posterior programme activity, while in the E5.5 epiblast the *Wt1*-responsive target programme defined experimentally *in vitro* is preferentially active in *Wt1*-high cells. Among E5.5 epiblast cells with informative *Wt1* splice reads, predominant KTS-representation aligns with increased anterior neural stem cell module expression, linking KTS splice status to anterior-associated transcriptional bias *in vivo*. Although these embryo analyses remain correlative, the concordance between splice-associated transcriptional shifts *in vivo* and isoform-dependent transcriptional programmes observed in stem cell models suggests that endogenous *Wt1* splice composition may already bias lineage programme engagement in the epiblast prior to gastrulation. By contrast, the posterior/mesoderm-associated activity of *Wt1* KTS+ is supported primarily by the *in vitro* differentiation experiments and is not equivalently demonstrated *in vivo during peri-implantation development*.

Spatially resolved analysis provides additional context for this association. At E5.5, when anterior-posterior organization is not yet clearly resolved within the epiblast, higher *Wt1* expression was associated with the side of the embryo showing stronger expression of *Dkk1* and *Cer1,* markers associated with the emerging AVE. Moreover, among E5.5 epiblast regions containing informative KTS splice-junction fragments, a higher fraction of KTS-transcripts was associated with a stronger anterior-biased transcriptional state. The temporal relationship between this early *Wt1* asymmetry and AVE-dependent patterning remains unresolved.

*Wt1* induction in differentiating naïve human ESCs reproduced the same regulatory logic observed in mouse. Although *Wt1* peak expression during the formative transition is low and evidence for expression at the in vivo post implantation epiblast stage in human embryos remains limited, the conserved transcriptional responses observed in human pluripotent cells indicate that the regulatory capacity of *Wt1* is preserved and can be engaged during early human development. This raises the possibility that species-specific differences in the temporal deployment of *Wt1* contribute to developmental pacing, with *Wt1* acting to accelerate formative progression in mouse, whereas its role in human development may be reduced or differently timed.

*Wt1 i*s widely implicated in cancer: approximately 20% of Wilms tumors carry *Wt1* mutations, whereas *Wt1* expression is elevated in many other malignancies, including acute myeloid leukemia^49,50^. This context dependence suggests that both loss and ectopic *Wt1* activity can contribute to tumorigenesis^23^. Our findings provide a developmental basis for this behaviour, showing that WT1 engages active regulatory elements and biases gene regulatory programmes that shape lineage competence rather than acting as a classical lineage determinant. Loss of *Wt1* function could therefore impair appropriate lineage programme engagement, whereas ectopic or sustained expression could redirect transcriptional states toward inappropriate fates. Such lineage mis-specification and transcriptional plasticity are characteristic of malignant progression^51^, suggesting that the recurrent involvement of *Wt1* in cancer may reflect pathological redeployment of its developmental lineage-regulatory activity. Although the name-giving Wilms tumour originates from kidney development and *Wt1* has historically been linked to renal lineages, several observations indicate that *Wt1* activity is not restricted to mesodermal contexts. *Wt1* expression and *WT1*-containing oncogenic fusions have been detected in human tumours with neural or neuroectodermal characteristics^52,53^ such as glioblastoma and retinoblastoma, and the EWS–WT1 fusion oncoprotein has been shown to activate neuronal transcriptional programmes^54^. Although *Wt1* splice isoforms have been reported in several tumor types, their functional significance in cancer remains poorly understood. The lineage-biasing activity we observe for *Wt1* isoforms in pluripotent cells raises the possibility that differential *Wt1* isoform activity could similarly influence lineage-associated transcriptional states in tumors, depending on the surrounding regulatory environment.

*Wt1* therefore emerges as an early regulator acting during formative pluripotency, a developmental window in which the epiblast acquires responsiveness to lineage-inducing cues. The transient expression of *Wt1* in the E5.5 epiblast, its association with emerging anterior transcriptional asymmetry, and the concordance between isoform-dependent transcriptional biases in stem cell models and splice-associated transcriptional differences in the embryo together indicate that *Wt1* activity shapes how lineage programmes are engaged during formative pluripotency, prior to overt lineage segregation. More broadly, these observations support a model in which lineage-associated transcriptional programmes diverge within transitional pluripotent states, rather than emerging only at discrete developmental transitions. This suggests that regulated lineage bias is an intrinsic feature of formative pluripotency that emerges before overt germ layer commitment.

## Acknowledgements

We thank Kitti Dóra Csályi, Thomas Sauer and Endre Kiss from the Max Perutz Laboratories FACS Facility for expert support. Next generation sequencing was performed at the Vienna Biocenter Core Facilities (VBCF). We thank Lejla Zijadic and Katherina Tavernini for experimental support. We thank all members of the Leeb and Buecker labs for critical comments and technical support, and Kristina Stapornwongkul and Florian Halbritter critical comments and helpful suggestions. This research was funded in whole, or in part, by the Austrian Science Fund (FWF; dois: 10.55776/I5958, 10.55776/P35637 and 10.55776/PAT8065924). For open access purposes, the author has applied a CC BY public copyright license to any author accepted manuscript version arising from this submission. L.M.C.A. is a Max Perutz fellowship awardee. M.H. and M.P. are members of the FWF-funded doctoral programme ‘Signalling Molecules in Cellular Homeostasis’ (SMICH; 10.55776/W1261). M.L. is a faculty member and speaker of SMICH and a Wissenschafts-Forschungs-und Technologiefonds (WWTF) Vienna Research Group Leader (VRG14-006).

## Data Availability

The RNA-seq data generated in this study have been deposited in the Gene Expression Omnibus (GEO) under accession code GSE326071, and the ChIP-seq data under accession code GSE326095. All other relevant data supporting the key findings of this study are available within the article and its Supplementary Information files.

## MATERIALS AND METHODS

### Cell culture

mESCs were maintained on tissue culture plates coated with gelatin (Sigma-Aldrich, G1890) at 37 °C in a humidified incubator with 5% CO₂. Cells were grown in high-glucose DMEM (Sigma-Aldrich, D5671) supplemented with 10% batch-tested FBS (Sigma-Aldrich, F7524, or Biowest, S1600), 2 mM L-glutamine (Sigma-Aldrich, G7513), 0.1 mM non-essential amino acids (Sigma-Aldrich, M7145), 1 mM sodium pyruvate (Sigma-Aldrich, S8636), 10 µg/ml penicillin–streptomycin (Sigma-Aldrich, P4333), 0.05 mM 2-mercaptoethanol (Gibco, 31350-010), 10 ng/ml LIF (batch-tested, produced in-house), and the MEK and GSK3 inhibitors PD0325901 (1.0 µM) and CHIR99021 (3 µM). This medium is referred to as ES DMEM-2i. Cells were passaged every 2 days using trypsin–EDTA and routinely tested negative for mycoplasma contamination.

Naïve SHEF6 hESCs were maintained on mitotically inactivated mouse embryonic fibroblast (MEF) feeder cells in PXGL medium at 37 °C in a humidified incubator. PXGL consisted of N2B27 supplemented with 1 µM PD0325901, 2 µM XAV939, 2 µM Gö6983 and 10 ng/ml human LIF. For differentiation, naïve SHEF6 cells were dissociated with Accutase and seeded onto Geltrex-coated plates in N2B27 medium supplemented with 10 µM Y-27632 at plating. From the following day, cells were cultured in N2B27 without Y-27632. *Wt1* KTS- and *Wt1* KTS+ forced-expression experiments were performed under these differentiation conditions and cells were harvested at the indicated time points for transcriptomic analysis.

### Cell line generation

For CRISPRa experiments, the piggyBac vector PB-TRE-dCas9-VPR (Addgene plasmid #63800) was introduced into Rex1-GFP mESCs. Transfected cells were selected in ES DMEM-2i with hygromycin. Single-cell clones were isolated, tested for doxycycline-inducible dCas9-VPR expression, and one representative clone was expanded and used for all CRISPRa experiments.

Knockout (KO) cell lines were generated in the RC9 background, which carries an EF1α-driven Cas9 cassette targeted to the Rosa26 locus in Rex1-GFP mESCs (RC9 cells, previously described in Li et al.^7^. Cells (10^5^) were reverse transfected in gelatin-coated six-well plates in ES DMEM-2i using Lipofectamine 2000 (Fischer Scientific 11668-027). Two gRNAs were used to knock out *Wt1* (gRNA1: TTCAAACACGAGGACCCCAT, gRNA2: AATCTCCTTCCGCTCATCGT). These gRNAs were cloned into a pB-hU6-gRNA-BFP vector. BFP+ cells were sorted and plated at low density in 6 cm TC treated culture dishes (Starlab CC7682-3359) and grown for 6 days prior to colony picking and expansion of knockout clones and genotyping by PCR.

For precocious *Wt1* expression, constructs encoding the four major splice isoforms, *Wt1* KTS-/1 (-Ex5/-KTS), *Wt1* KTS-/2 (+Ex5/-KTS), *Wt1* KTS+/1 (+Ex5/+KTS) and *Wt1* KTS+/2 (-Ex5/+KTS), were cloned into a pB-TetOn-3xFLAG-Empty-PolyA-puromycin vector. *Wt1* was amplified from RNA using Superscript III with the following primers designed for cloning: forward 5’-AGAGGCGCGCCCTGGACTTCCTCCTGTCGCAG-3’, reverse: 5’-ACTCCAGCTGGCGCTTTGACTCGAGTAG-3’.

The constructs for doxycycline-inducible forced expression were delivered into the Rex1::GFPd2 mESCs. Precocious expression of Otx2 was achieved by delivering a vector pB-TetOn-Otx2-Puromycin obtained from Christa Buecker (Max Perutz Labs).

WT1 Knockout cell lines and forced/precocious expression were corroborated by western blot.

### Monolayer differentiation

To induce exit from naïve pluripotency, mESCs grown in ES DMEM-2i were dissociated to single cells and replated on gelatin-coated plates at 1 × 10⁴ cells/cm² in N2B27-2i. N2B27 consisted of a 1:1 mixture of DMEM/F12 (Gibco, 21331020) and Neurobasal medium (Gibco, 21103049) supplemented with homemade N2, B27 Serum-Free Supplement (Gibco, 17504-044), 2 mM L-glutamine (Sigma-Aldrich, G7513), 0.1 mM NEAA (Sigma-Aldrich, M7145), 10 µg/ml penicillin–streptomycin (Sigma-Aldrich, P4333) and 55 µM β-mercaptoethanol (Fisher Scientific, 21985-023). N2B27-2i was obtained by adding 1 µM PD0325901 and 3 µM CHIR99021. After 16–20 h, cultures were washed once with pre-warmed PBS and the medium was changed to N2B27 (referred to as N medium). The time of this medium switch was defined as N0. Subsequent time points in differentiation conditions are indicated as Nx, where x denotes hours after 2i withdrawal.

### EB differentiation assay

EBs were generated following a standard hanging drop protocol^55^. EBs were formed using 150 x 20 mm dishes (CytoOne, CC7682-3614) using ESC culture medium without LIF and 2i (200 cells/ 20 µL drop). After 2 days, EBs were collected in 9 cm non-TC treated plates (Thermo Scientific, 11389273); the medium was changed after 2 days. RNA was extracted using the KingFisher Flex Purification System (Thermo Fisher) with the High-Performance RNA Isolation kit from the Molecular Tools Shop (Vienna, Biocenter).

### Differentiation into NSC

NSCs used as a reference for NESTIN-positive cells were generated by following standard protocol^56^ consisting of plating cells into N2B27 for 7 days in a gelatin-coated flask (CytoOne, CC7682-4875). Then the cells were split using trypsin–EDTA and plated into a non-coated flask containing N2B27 supplemented with EGF (Preprotech, 315-09-100UG) and FGF (Preprotech, 315-09-100UG) (10 ng/ml), referred to as NSC medium. Three days later, neurospheres formed from floating cells were plated into a gelatin-coated flask. Cells were split when reaching 50% confluency and later, every 2 days in NSC medium using accutase (Thermo Scientific, A1110501). For programming of ESCs toward NSCs by transient *Wt1* KTS-forced expression, doxycycline was applied for the initial 48 h, followed by washout and a further 24 h in doxycycline-free NSC medium before analysis.

### Flow cytometry analysis

For flow cytometric analysis of Rex1::GFPd2-derived cell lines, cells were harvested at the indicated time points by dissociation with 0.25% trypsin-EDTA and neutralised in ES DMEM. Cell suspensions were passed through 40 µm strainers to remove aggregates and kept on ice until acquisition. Forward and side scatter parameters and GFP fluorescence were recorded on an LSR Fortessa flow cytometer (BD Biosciences). Debris and doublets were excluded by sequential gating on FSC/SSC and FSC-A/FSC-H. Data were analysed using FlowJo (BD Biosciences). Rex1-GFP levels were quantified either as the percentage of GFP⁺/⁻ cells, defined by gates set on control samples.

### Immunofluorescence

For immunofluorescence experiments, mESCs were plated at 1 × 10⁴ cells/cm² on fibronectin-coated multiwell glass slides (Marienfeld Superior, 1216330) in N2i for 16–20 h. The medium was either changed to N2B27 to start differentiation or replaced with fresh N2i for naïve controls. Cells were fixed at the indicated stages (2i, N24, N42) by washing once with PBS and incubating for 15 min at room temperature in 4% paraformaldehyde in PBS. Fixed cells were rinsed in PBS, permeabilized for 10 min in PBS with 0.1% Triton X-100, and then incubated for 60 min in PBS containing 0.1% Tween-20 and 5% BSA to block non-specific binding.

Primary antibody was applied overnight at 4 °C. The next day, cells were washed three times in PBS with 0.1% Tween-20 and incubated for 1 h at room temperature in the dark with a far-red Alexa Fluor– conjugated anti-rabbit secondary antibody (1:500), together with DAPI (1 µg/ml) to label nuclei. After three additional washes in PBS, slides were kept in PBS at 4 °C until imaging. Both primary and secondary antibodies were diluted in blocking buffer and used 1:1000. WT1 was detected using a rabbit anti-WT1 antibody (Abcam, ab89901, 1:100), NESTIN (Merck, MAB353, 1:100).

Images were acquired on a Zeiss Axio Observer Z1 microscope using a 40× objective, with identical acquisition settings applied to all conditions within a given experiment unless indicated otherwise. Image analysis was performed in Fiji/ImageJ.

### Immunoblotting

For immunoblotting, cells grown in ES DMEM-2i or N2B27 were rinsed once with ice-cold PBS and lysed directly on the plate in RIPA buffer supplemented with protease and phosphatase inhibitors. Lysates were incubated on ice for 20 min and then clarified by centrifugation at 13,000 rpm for 30 min at 4 °C. Protein concentration in the supernatant was measured using a colorimetric assay (Bradford or BCA) according to the manufacturer’s instructions. Equal amounts of protein (typically 10–20 µg per lane) were mixed with Laemmli sample buffer containing β-mercaptoethanol, heated at 95 °C for 5 min and separated by SDS–PAGE on 8–12% polyacrylamide gels. Proteins were transferred to nitrocellulose membranes and blocked for 1 h at room temperature in 5% milk in PBS with 0.1% Tween-20 (PBS-T).

Membranes were incubated with primary antibodies diluted in blocking buffer for 1–2 h at room temperature or overnight at 4 °C. WT1 was detected using rabbit anti-WT1 (Abcam, ab89901, 1:1000); VINCULIN was used as a loading control and detected with an anti-VINCULIN antibody (Santa Cruz Biotechnology, sc-25336, 1:5000). After three washes in PBS-T, membranes were incubated for 1 h at room temperature with HRP-conjugated secondary antibodies, washed again and developed using chemiluminescent substrate. Western blots were imaged and analyzed using an Invitrogen™ iBright™ FL1500 Imaging System (Thermo Fisher Scientific) and a ChemiDoc™ Touch Imaging System (Bio-Rad) and images processed with ImageJ/Fiji and iBright Analysis Software V5.

### CRISPRa screen and gRNA enrichment analysis

A targeted CRISPRa screen was performed to identify formative-induced genes (FIGs) whose activation promotes exit from naïve pluripotency under naïve-stabilizing conditions. Rex1::GFPd2 mESCs carrying a doxycycline-inducible dCas9–VPR transgene were transduced with a pooled sgRNA library comprising 1,298 gRNAs. The library contained 1,180 gRNAs targeting promoter-proximal regions of FIGs, with 5–10 independent gRNAs per gene and included sgRNAs targeting Otx2 as a reference. Control guides comprised 59 sgRNAs targeting non-existent sequences and 59 sgRNAs targeting non-annotated genomic regions. Following enrichment of sgRNA-positive cells, CRISPRa activity was induced by doxycycline while cells were maintained in 2i conditions. After 60 h of induction, cells were dissociated into single-cell suspensions and sorted by flow cytometry into GFP-high (Rex1-GFP⁺) and GFP-low/negative (Rex1-GFP⁻) fractions. Genomic DNA was isolated from each sorted population, and sgRNA cassettes were PCR-amplified using Q5 High-Fidelity DNA Polymerase (NEB) with Illumina-compatible primers. Amplicons were sequenced at the Vienna BioCenter Core Facilities (VBCF) NGS facility on an Illumina NovaSeq SP SR100 XP run to quantify guide abundance. Differential sgRNA enrichment between GFP⁻ and GFP⁺ fractions was assessed using MAGeCK^57^ implemented through the Galaxy platform^58^ ,and replicate concordance was evaluated by comparing sgRNA count profiles between samples.

### Chromatin immunoprecipitation (ChIP)

WT1 and FLAG ChIP were performed based on methods previously described^59^. Cells were plated on gelatin-coated 15 cm dishes, washed with PBS and crosslinked directly on the plate in PBS containing 1% formaldehyde for 10 min at room temperature, followed by quenching with 0.125 M glycine for 10 min. Cells were washed twice with cold PBS, harvested by scraping in ice-cold PBS containing 0.01% Triton X-100, pelleted, washed once in cold PBS, flash-frozen in liquid nitrogen and stored at -80 °C. Frozen pellets were thawed on ice, and lysis buffers were supplemented with protease inhibitors immediately before use. Nuclei were extracted by sequential lysis in LB1 (50 mM HEPES-KOH pH 7.5, 140 mM NaCl, 1mM EDTA, 10% glycerol, 0.5% NP-40, 0.25% Triton X-100) for 10 min at 4 °C with rotation, followed by centrifugation (1,350 x g, 5 min, 4 °C), and LB2 (10 mM Tris-HCl pH 8.0, 200 mM NaCl, 1mM EDTA, 0.5 mM EGTA) for 10 min at room temperature with rotation. After a second centrifugation step (1,350 xg, 5 min, 4 °C), pellets were resuspended in LB3 (10 mM Tris-HCl pH 8.0, 100 mM NaCl, 1mM EDTA, 0.5 mM EGTA, 0.1% sodium deoxycholate, 0.5% N-lauroylsarcosine), supplemented with sonication beads, and chromatin was fragmented by sonication for 12–15 cycles (30 s on/45 s off) with cooling. Lysates were cleared by centrifugation (16,000 x g, 10 min, 4 °C), and Triton X-100 was added to the supernatant to a final concentration of 1%. For each immunoprecipitation, 1 mL of chromatin was incubated with antibody overnight at 4 °C with rotation, and 10% input material was retained. Endogenous WT1 ChIP was performed using anti-WT1 (Abcam, ab89901; 12 μL per IP, corresponding to ∼2.6 μg), and FLAG ChIP was performed using anti-FLAG M2 (mouse; Sigma, F1804; 5 μg per IP). Immune complexes were captured using 100 μL Dynabeads Protein G per IP, which were washed three times in cold block solution (PBS containing 0.5% (w/v) BSA) before incubation with antibody-bound chromatin for 2-4 h at 4 °C with rotation. Beads were washed at least five times with cold RIPA wash buffer (50 mM HEPES-KOH pH 7.5, 500 mM LiCl, 1 mM EDTA, 1% NP-40, 0.7% sodium deoxycholate) and once with TE containing 50 mM NaCl. Chromatin was eluted in 210 μL elution buffer (50 mM Tris-HCl pH 8.0, 10 mM EDTA, 1% SDS) for 15 min at 65 °C with intermittent mixing, and inputs were adjusted to matched elution conditions. ChIP and input samples were reverse-crosslinked overnight at 65 °C, diluted 1:1 with TE buffer, treated with RNase A (0.2 mg/mL final; 2 h at 37 °C), and subsequently incubated with Proteinase K (0.2 mg/mL final; 30 min at 55 °C) after adjustment of CaCl_2_ to 5.25 mM. DNA was purified by phenol-chloroform extraction using Phase Lock Gel tubes, followed by ethanol precipitation using sodium acetate and glycogen as carrier, washed with 70% ethanol, air-dried and resuspended in water for downstream library preparation.

### ChIP-sequencing analysis

WT1 and FLAG ChIP-seq datasets were processed with nf-core/chipseq to generate coordinate-sorted BAM files aligned to the mouse reference genome (mm10). WT1 ChIP-seq reads were processed with nf-core/chipseq v2.1.0-g76e2382 (Nextflow v25.04.3; bowtie2 v2.5.2), whereas FLAG ChIP-seq reads were processed with nf-core/chipseq v2.0.0 (Nextflow v23.10.1; bowtie2 v2.4.4). For both datasets, peak calling was performed outside the nf-core pipeline using the resulting BAM files and MACS3 (v3.0.3) in narrowPeak mode with input DNA as control. For each condition/time point, biological replicate BAM files were pooled for peak calling to generate a single peak set per condition, using pooled input BAM files as control. Peak sets were filtered to remove peaks mapping to non-canonical chromosomes, including unplaced or unlocalized contigs (for example chrUn and other non-canonical contigs), and peaks overlapping the ENCODE/Boyle Lab mm10 blacklist (v2). To restrict analyses to high-confidence binding events, peaks were additionally filtered by MACS peak score, retaining peaks with score ≥100, based on empirical inspection in IGV indicating reduced reliability of lower-scoring peaks.

For WT1 ChIP-seq, ChIP was performed at N2i (n = 2), N24 (n = 4) and N42 (n = 4). Input DNA libraries (n = 4) were pooled and used as a common control for peak calling across stages. For each stage, replicate BAM files were pooled for peak calling to generate a single WT1 peak set per time point against the pooled input dataset.

For FLAG ChIP-seq, ChIP was performed for 3×FLAG-tagged WT1 KTS-under high induction (HE; 250 ng/mL doxycycline) in naïve-stabilizing conditions (2i HE) and in differentiation-permissive conditions (N16 HE). Each FLAG condition was represented by two biological ChIP replicates (n = 2 per condition), which were pooled for MACS3 peak calling to generate a single peak set per condition, using pooled input DNA as control.

For peak-to-gene assignment used in Fig. 2c, WT1 ChIP-seq peaks were annotated in R using ChIPseeker 1.42.1 against mm10 gene annotations, and peaks were associated to nearby genes according to the ChIPseeker peak annotation framework. For genomic feature classification shown in Supplementary Fig. 2b, a hierarchical annotation strategy was used. Peaks within ±1 kb of annotated transcription start sites were first classified as promoter-proximal. Promoter-proximal peaks were then excluded, and distal peaks were intersected with published ESC- and EpiLC-enhancer coordinates (including an ESC/EpiLC shared enhancer set) from the Bücker laboratory dataset using bedtools, and assigned based on overlap with ESC, EpiLC, or shared enhancers. Peaks not overlapping any enhancer category were subsequently annotated with ChIPseeker to classify remaining peaks into intronic and other/intergenic categories.

For global comparisons across datasets, the genomic background was defined as 155266 regions, corresponding to merged ATAC-seq peaks and enhancer peaks as used in Santini et al.^60^. For feature-level comparisons in Supplementary Figure S3b, background estimates were referenced to mm10 (GRCm38.p6) using the GENCODE M25 annotation. Promoters were approximated from the number of annotated mouse protein-coding genes (21859). Enhancers were defined using the curated ESC/EpiLC enhancer framework, with TSS-overlapping regions excluded, and classified as ESC-only (9557), EpiLC-only (23554), or shared (11021)^59^. Introns were considered an approximate transcript-associated compartment (∼101,000 regions), estimated from GENCODE M25 annotation-derived protein-coding gene content and representative transcript-based intron number per gene.

### Tier-based analysis of FLAG–WT1 KTS– HE

Tier-based analyses were performed on the subset of 3xFLAG–WT1 KTS– (Exon5–, KTS–) ChIP–seq peaks that overlapped endogenous WT1 peaks at N42. Peak overlaps between FLAG and WT1 datasets were computed using bedtools (v2.30.0) after coordinate sorting (bedtools sort), and overlapping peaks were defined using bedtools intersect. For tiering, the shared peak set was ranked by FLAG ChIP signal intensity using the corresponding FLAG BigWig track for the condition of interest (2i + doxycycline or N16 + doxycycline). Signal was quantified per peak from the BigWig and peaks were ordered by decreasing FLAG signal; the ranked list was then partitioned into five equal-sized bins to define Tier 1 (highest FLAG signal) through Tier 5 (lowest FLAG signal). Peaks detected in the FLAG ChIP–seq datasets that did not overlap endogenous WT1 peaks at N42 were classified as “FLAG-only” and analysed as a separate category. Peak-to-gene annotation for each tier and for the FLAG-only category was performed in R using ChIPSeeker (v1.34.1), and gene set enrichment analyses were carried out separately for each tier using the resulting tier-associated gene lists.

### Long-read amplicon sequencing for WT1 isoform analysis

Total RNA was extracted using a KingFisher-based magnetic bead workflow (Thermo Fisher Scientific) and reverse-transcribed using SuperScript™ III Reverse Transcriptase (Thermo Fisher Scientific, cat. no. 18080093). WT1 transcripts were amplified from cDNA by PCR using isoform-discriminating primer pairs spanning informative exon-exon junctions (same primers used to clone WT1 into expression vector). Amplicons were submitted for long-read sequencing (Plasmidsaurus, Amplicon PCR Premium). N42 samples were sequenced in technical duplicate. FASTQ files returned by the provider were used for downstream isoform discovery and quantification.

Reads were aligned to the mouse reference genome (mm10) using minimap2 in splice-aware mode. Alignments were generated in SAM format, converted to coordinate-sorted BAM files using SAMtools, and filtered to retain only primary alignments for isoform analysis. Transcript isoform discovery and correction were performed using FLAIR (Full-Length Alternative Isoform analysis of RNA). Briefly, aligned reads were converted to BED12 format, splice junctions were corrected using flair correct with the mm10 reference genome and a curated gene annotation (GTF), and corrected reads were collapsed into non-redundant transcript models using flair collapse. Isoform-level expression was quantified using flair quantify using default read-assignment criteria based on alignment quality, fraction of read aligned, and fraction of transcript covered. To maximize sensitivity in distinguishing closely related isoforms, the minimum MAPQ threshold was set to 0 and only primary alignments were considered. Transcript-level count matrices were generated for downstream comparative analyses.

Protein-coding WT1-bound genes annotated at N42 were subjected to over-representation analysis using the Enrichr platform^61^ with the ChEA 2022 and ENCODE and ChEA Consensus TFs from ChIP-X gene-set libraries^62,63^.

### Comparison with ChIP-seq datasets

Public ChIP–seq and ATAC–seq datasets were compiled and harmonized for comparison with WT1 by ensuring a consistent mm10 reference throughout. Datasets previously reported in literature for OCT4 ESC, OCT4 EpiLC, OTX2, p300 ESC, p300 EpiLC^10^ were obtained in BED (peaks) format (and BigWig tracks when available) and processed for downstream overlap/visualization. Datasets previously used and processed in Santini et al.^60^ for FOXO1 2i, FOXO1 N25, ATAC 2i, ATAC N25 were likewise obtained as BED peak files (and BigWigs where applicable) and processed identically. ESRRB datasets from published datasets^25^ (ESRRB 48h) were retrieved in peak/track formats consistent with mm10 and processed in the same manner. NANOG and SOX2 datasets^64^ were obtained as BED peaks and BigWigs when available and processed for downstream analyses. For the TCF7L1/TCF3 dataset^65^ used in this analysis, aligned reads were not available as standard BAMs, reads were reconstructed by mapping to mm10 (bowtie2), generating coordinate-sorted/indexed BAMs (samtools), producing BigWigs (bedtools genomecov and UCSC bedGraphToBigWig), and calling peaks with MACS3 (v3.0.3). For remaining datasets involving additional TFs and Polycomb-associated factors SUZ12^66^ and PCGF2^67^: peak calling was performed with MACS3 (v3.0.3) from BAM files when required (narrowPeak mode for TF-like datasets; broad peak calling where appropriate for broader chromatin-associated signals), generating BED peak sets for downstream overlap and annotation analyses. Across all datasets, peak BED files were standardized by removing peaks on non-canonical contigs (including chrUn/unplaced/unlocalized contigs), excluding regions overlapping the ENCODE/Boyle Lab mm10 blacklist (v2), and applying an IGV-guided peak-confidence filter (retaining peaks with MACS peak score ≥100 or ≥50 for broader/non-motif-like datasets after manual inspection on IGV browser, yielding final mm10 BED peak sets and representative BigWig tracks for visualization.

In addition, an H3K27ac reference dataset^68^ was incorporated for comparative analyses; peaks were generated/used in mm10 and, when peak calling was required, MACS3 (v3.0.3) broad peak calling was applied before blacklist/non-canonical contig removal and IGV-guided peak-score filtering as described above.

### Single cell data analysis from differentiation time course

Wild-type RC9 mESCs were plated at 10,000 cells/cm² in N2B27-2i medium. Differentiation was initiated the following day by washing cells once with PBS and switching to N2B27 medium. Cells were collected after 12 h, 24 h, or 48 h of differentiation. For each time point, two biological replicates were generated from independent cultures (n = 2).

Cells were harvested by trypsinization, resuspended in ice-cold PBS supplemented with 0.04% BSA (Fisher Scientific, 15260037), and adjusted to 1,000 cells/µL. Downstream processing was performed by the VBCF NGS core facility. Cell viability was assessed by acridine orange staining and exceeded 80% for all samples. For each sample, 100,000 cells were immediately fixed in 4% formaldehyde using the “Fixation of Cells & Nuclei for Chromium Fixed RNA Profiling” protocol (10x Genomics) with the Single Cell Fixed RNA Sample Preparation Kit (10x Genomics, PN-1000414). Fixed cells were individually barcoded using whole-transcriptome probe pairs with the Chromium Fixed RNA Kit (Mouse Transcriptome; 10x Genomics, PN-1000497) and subsequently pooled for library preparation and sequencing. Libraries were sequenced on an Illumina NovaSeq 1B in paired-end 150 bp mode. Approximately 8,000 cells were sequenced per sample. Primary processing of sequencing data was performed using Cell Ranger.

For downstream analysis, data were processed in R using Seurat. Features detected in at least 10 cells were retained, and cells with fewer than 200 detected features were excluded. Data were log-normalized, and replicate datasets were merged for visualization and plotting.

For Fig. 4c, anterior and posterior module scores were computed in N48 single-cell RNA-seq data from predefined gene sets (anterior: *Otx2*, *Zic1*, *Zic3*, *Pou3f1*, *Sox2*, *Sox3*, *Six3* and *Lhx2*; posterior: *T*, *Eomes*, *Mixl1*, *Mesp1*, *Foxa2*, *Fgf8*, *Wnt3*, *Snai1* and *Lefty2*). For analyses stratified by *Otx2* expression, *Otx2* itself was excluded from the anterior module to avoid circularity. Cells were assigned to high- and low-expression groups for *Wt1*, *Otx2* and *Nanog* using log-normalized expression values, with values >0.5 classified as high and values ≤0.5 classified as low. Differences in mean module scores between groups were evaluated by permutation analysis using 50,000 random label shuffles.

### Single-cell reference integration

A reference atlas of inner cell mass (ICM) and epiblast (Epi) cells spanning embryonic days E3.5–E6.5 was assembled from five publicly available single-cell RNA-seq datasets^18,26,34–37^. Datasets were batch-corrected and integrated using FastMNN implemented in the batchelor package (v1.24.0), yielding a corrected low-dimensional representation and expression matrix for downstream analyses. The integrated reference was analysed using the standard Seurat workflow (v5.3.0). As query, ComBat-corrected bulk RNA-seq expression profiles were converted into a Seurat object, treating each bulk sample as a single query observation (pseudo-cell) after LogNormalize transformation. Anchors between reference and query were identified using FindTransferAnchors (normalization.method = “LogNormalize”, reference.reduction = “pca”, dims = 1:50). Reference-derived developmental labels were assigned to query observations using TransferData. To embed bulk query observations into the reference latent space, IntegrateEmbeddings was performed using the computed anchors with k.weight = 5, and reference and query objects were merged for visualization of query mapping relative to the integrated *in vivo* reference.

### RNA-sequencing analysis

Bulk RNA-seq was performed using two complementary workflows depending on the experiment. For mouse mESC experiments (monolayer differentiation and embryoid body conditions, including KO and WT1 forced-expression datasets), total RNA was submitted to the Vienna BioCenter (VBCF) NGS facility for 3′-end, strand-specific library preparation using the Lexogen QuantSeq kit and sequencing on an Illumina NovaSeqX platform (NovaSeqX 10B XP flowcell; single-end 100 cycles for monolayer HE/FE datasets; paired-end 150 cycles for KO/EB and FE-in-EB datasets). Unless indicated otherwise, experiments were performed with two biological replicates; for HE expression, two technical replicates per biological replicate were sequenced; for FE conditions and KO experiments, two independent clonal lines were analysed with two replicates per clone. Fastq files generated by the VBCF facility were processed using nf-core/rnaseq (v3.21.0) against the mouse reference genome (mm10; GENCODE annotation) using STAR together with Salmon quantification in pseudo-alignment mode. UMI-aware processing was enabled in nf-core/rnaseq using UMI-tools with regex-based UMI extraction (pattern matching the QuantSeq/UMI read structure; exact pattern as in the analysis pipeline command). Differential expression analyses for mouse mESC datasets, including FE and HE conditions in monolayer differentiation, and FE experiments in EBs (*Wt1* KTS+/1 and *Wt1* KTS-/1), and *Wt1* knockout samples in monolayer conditions, were performed in R using DESeq2. For knockout analyses, Y-linked genes were excluded prior to differential expression testing. Differentially expressed genes were defined based on the indicated thresholds for adjusted p value and log2 fold change used in each analysis.

For *Wt1* KTS+/2 and *Wt1* KTS-/2 FE samples in EBs, and for human WT1 FE experiments in SHEF6 cells, 3′-end counting RNA-seq (DGE-style) was performed by Plasmidsaurus on the Illumina platform using UMI-labeled cDNA generation and UMI-based deduplication. FASTQ quality was assessed with FastQC, followed by read quality filtering with fastp (including poly-X and 3′ tail trimming, minimum Phred score filtering, and minimum read-length filtering). Filtered reads were aligned to their corresponding reference genome using STAR, coordinate-sorted with samtools, and deduplicated using UMI-based collapsing (UMIcollapse). Alignment quality, strand specificity, and genomic feature distributions were assessed using RSeQC and Qualimap, and summary metrics were aggregated with MultiQC. Gene-level counts were generated with featureCounts and annotated using metadata from the provider’s gene annotation (GTF). Downstream analyses used the Plasmidsaurus-provided gene-level count tables for differential expression analysis as required for the study.

Enrichment analyses were performed using CellMarker 2024^69^, Reactome Pathways^70^ using the Enrichr platform ^61^. DEGs were also subjected to Gene Ontology Biological Process over-representation analysis using clusterProfiler and Gene Ontology annotations^71,72^.

To assess WT1 target behaviour under wild-type differentiation conditions for the analysis shown in Fig. 2f, the observed mean log_2_FC of WT1 target genes was compared with a null distribution generated by repeatedly sampling random sets of non-target genes matched to the target set size. Sampling was performed without replacement for 20,000 iterations, and the mean log2FC was calculated for each resampled set. Empirical p values were defined as the fraction of null means that were at least as extreme as the observed WT1 target mean. For the RNA-seq analysis shown in Fig. 5c, transcriptional changes are displayed as a log2FC-based heatmap, with genes selected from manually curated lists of lineage markers and cross-checked against the CellMarker 2024^69^.

### Analysis of *in vivo* RNA-seq data

Raw sequencing reads were downloaded in FASTQ format from the Sequence Read Archive using SRA Toolkit (v3.3.0), excluding technical reads. Reads were aligned to the mouse reference genome (mm10) using STAR (v2.7.11b) with default parameters for spliced alignment. Gene expression quantification was performed using Salmon (v1.10.3).

Samples were assigned to groups based on the embryonic day and tissue annotations reported in the original study. For epiblast cells from embryonic day E5.5 and E6.5, WT1 splice junction usage corresponding to KTS+/- isoforms was determined from STAR-generated SJ.out.tab files. Junction-supporting reads were annotated according to the following mm10 coordinates KTS+ (chr2:105170019-105172222), and KTS-(chr2:105170010-105172222). Cells were classified according to the predominant junction-supporting reads detected per cell. Because only E5.5 epiblast cells showed sufficient coverage to enable confident assignment of both splice events, subsequent analyses were restricted to this population.

Salmon-derived count matrices were imported into Seurat (v5.4.0) and normalized using the standard Seurat workflow. For comparisons of anterior and posterior gene-module activity, per-cell module scores were calculated using the Seurat AddModuleScore function. The anterior module comprised *Otx2, Zic1, Zic3, Pou3f1, Sox2, Sox3, Six3* and *Lhx2*, whereas the posterior module comprised *T, Eomes, Mixl1, Mesp1, Foxa2, Fgf8, Wnt3, Snai1* and *Lefty2*. For analyses stratified by *Otx2* expression, *Otx2* itself was excluded from the anterior module to avoid circularity. AddModuleScore calculates relative module activity by comparing the average expression of genes in the module with matched control features; therefore, scores below zero indicate lower relative module activity and do not imply absence of expression of the corresponding genes.

For analyses stratified by *Wt1, Otx2* and *Nanog* expression, cells were assigned to high- and low-expression groups using log-normalized expression values, with values >0.5 classified as high and values ≤0.5 classified as low. For comparisons of anterior and posterior gene-module activity in vivo (Fig. 7), the same predefined gene sets used in Fig. 4c were applied. Differences in mean module scores between groups were assessed by permutation analysis using 50,000 random label shuffles.

### Spatial analysis of *Wt1* KTS splice composition and anterior-posterior transcriptional bias

For the spatial analysis shown in Supplementary Fig. S7e, published GEO-seq FPKM expression matrices were analysed separately for each embryo. The anterior module comprised *Otx2, Zic1, Zic3, Pou3f1, Sox2, Sox3, Six3* and *Lhx2*, whereas the posterior module comprised *T, Eomes, Mixl1, Wnt3, Snai1* and *Lefty2*. FPKM values were transformed as log_2_(FPKM + 1), and each gene was subsequently z-scored across the spatial samples of the corresponding embryo. For each spatial sample, anterior and posterior module scores were calculated as the mean z-score of the genes in the respective module, and an anterior-minus-posterior score was calculated as the anterior module score minus the posterior module score.

*Wt1* KTS splice composition was quantified independently using strict fragment-level classification. The KTS-fraction was calculated as the number of informative fragments supporting KTS-divided by the total number of informative fragments supporting either KTS+ or KTS-. Samples lacking informative KTS fragments were excluded from the analysis. For the analysis presented in Supplementary Fig. S7e, only epiblast-derived spatial sections were retained.

## Declaration of generative AI and AI-assisted technologies in the writing process

During the preparation of this work the authors used ChatGPT in order to improve readability and language without changing scientific content. After using this tool, the authors carefully reviewed and edited the content as needed and take full responsibility for the content of the published article.

---

**Supplementary Figure 1.**
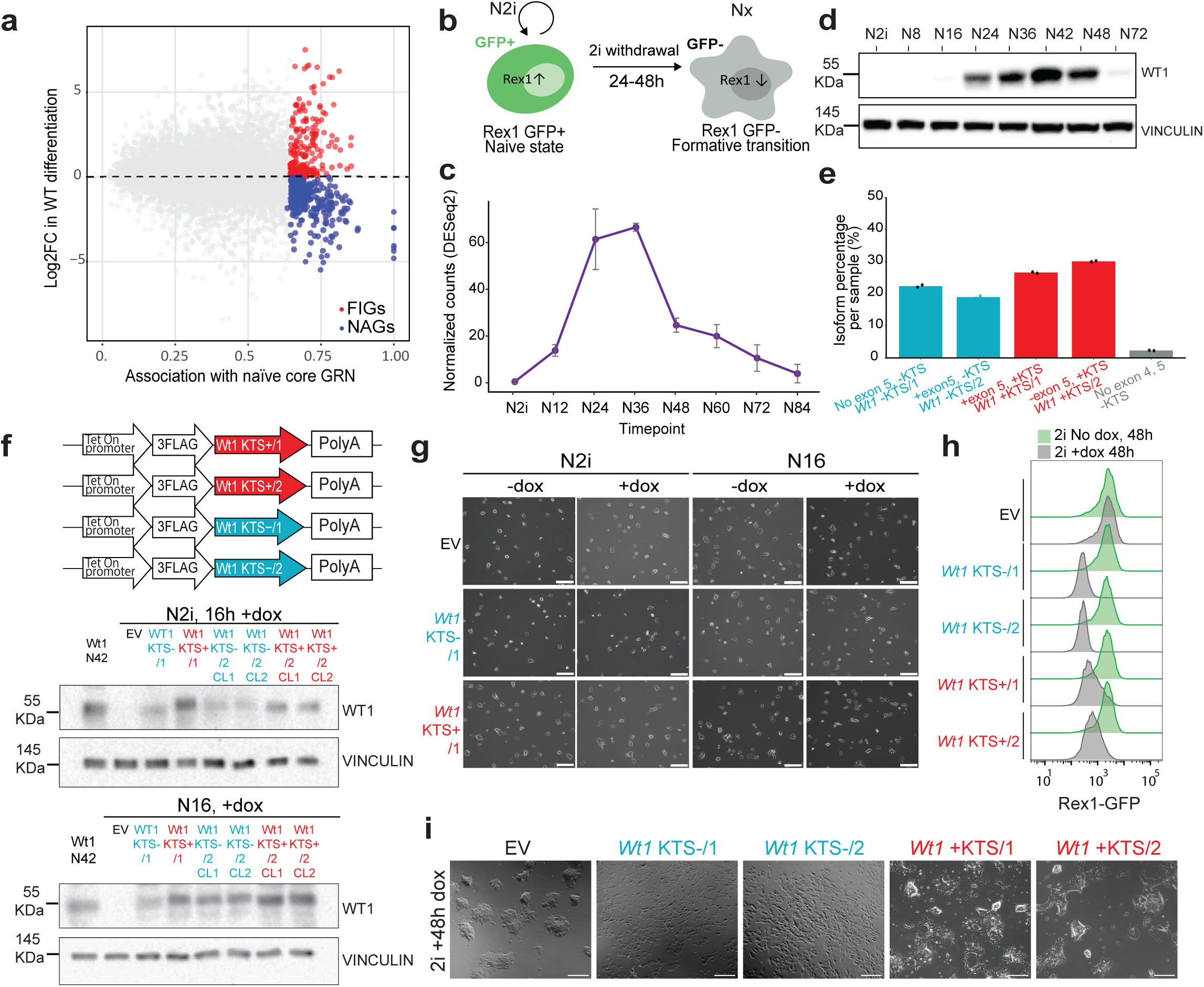
Precocious *Wt1* expression accelerates progression toward post-naïve epiblast identity a. Comparison of naïve network association (R²) and gene expression change during the naïve-to-formative transition. FIGs (red) and naive associated genes NAGs (blue) encompass genes that are highly anticorrelated and correlated to the naïve network, respectively. Modified from Lackner et al.^9^. b. Schematic of the experimental setup and Rex1::GFPd2 (Rex1-GFP) reporter readout during exit from the naïve state. Nx indicates x hours after 2i withdrawal in N2B27. One representative schematic is shown. c. Bulk RNA-seq time course showing transcriptional dynamics during exit from the naïve state. Data from a published naïve-to-formative RNA-seq time course^25^. d. Time course of WT1 protein expression during differentiation showing progressive upregulation and peak expression at N42. e. Quantification of endogenous *Wt1* splice-isoform abundance from cDNA at peak expression. Values are expressed as percentages. f. Schematic of the doxycycline-inducible forced-expression constructs used to generate the four *Wt1* splice isoforms, followed by western blot validation of doxycycline-induced *Wt1* expression in N2i and N16 conditions after 16 h of induction. EV, empty vector control. *Wt1* KTS-/1, *Wt1* KTS-/2, *Wt1* KTS+/1 and *Wt1* KTS+/2 are shown together with two independent clonal lines per construct (CL1 and CL2). Endogenous WT1 expression at N42 is shown as a reference. VINCULIN serves as a loading control. g. Representative bright-field images of empty vector control, *Wt1* KTS-/1 and *Wt1* KTS+/1 cells under N2i and N16 conditions following doxycycline induction. h. Representative Rex1-GFP flow cytometry profiles following 48 h doxycycline induction in N2i for empty vector control and the four WT1 splice isoforms. i. Representative bright-field images corresponding to the 48 h doxycycline induction experiment shown in (h), comparing empty vector control with *Wt1* KTS-/1, *Wt1* KTS-/2, *Wt1* KTS+/1 and *Wt1* KTS+/2 in N2i.

**Supplementary Figure 2.**
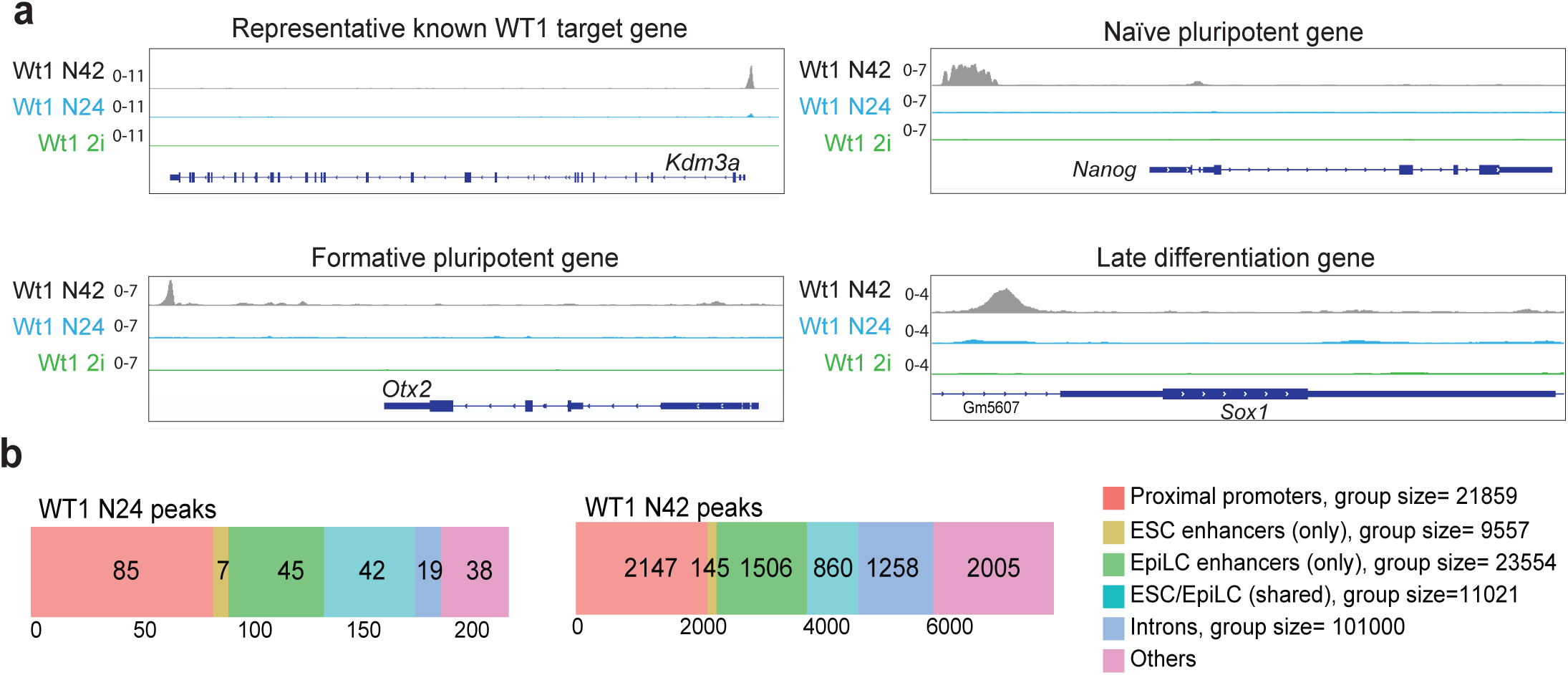
WT1 engages promoters and enhancers of developmental regulators in formative pluripotent cells a. Genome browser snapshots displaying WT1 ChIP-seq signal at representative loci in N2i (green), N24 (light blue), and N42 (gray). Shown genes include *Kdm3a* (as a positive-control locus used to benchmark ChIP performance), *Nanog* (naïve marker), *Otx2* (formative marker), and *Sox1* (post-implantation gene). b. Genomic feature distribution of WT1 peaks at N24 and N42. Peaks were annotated with ChIPseeker using ESC and EpiLC enhancer datasets. Group sizes represent the number of elements assigned to each genomic feature category.

**Supplementary Figure 3.**
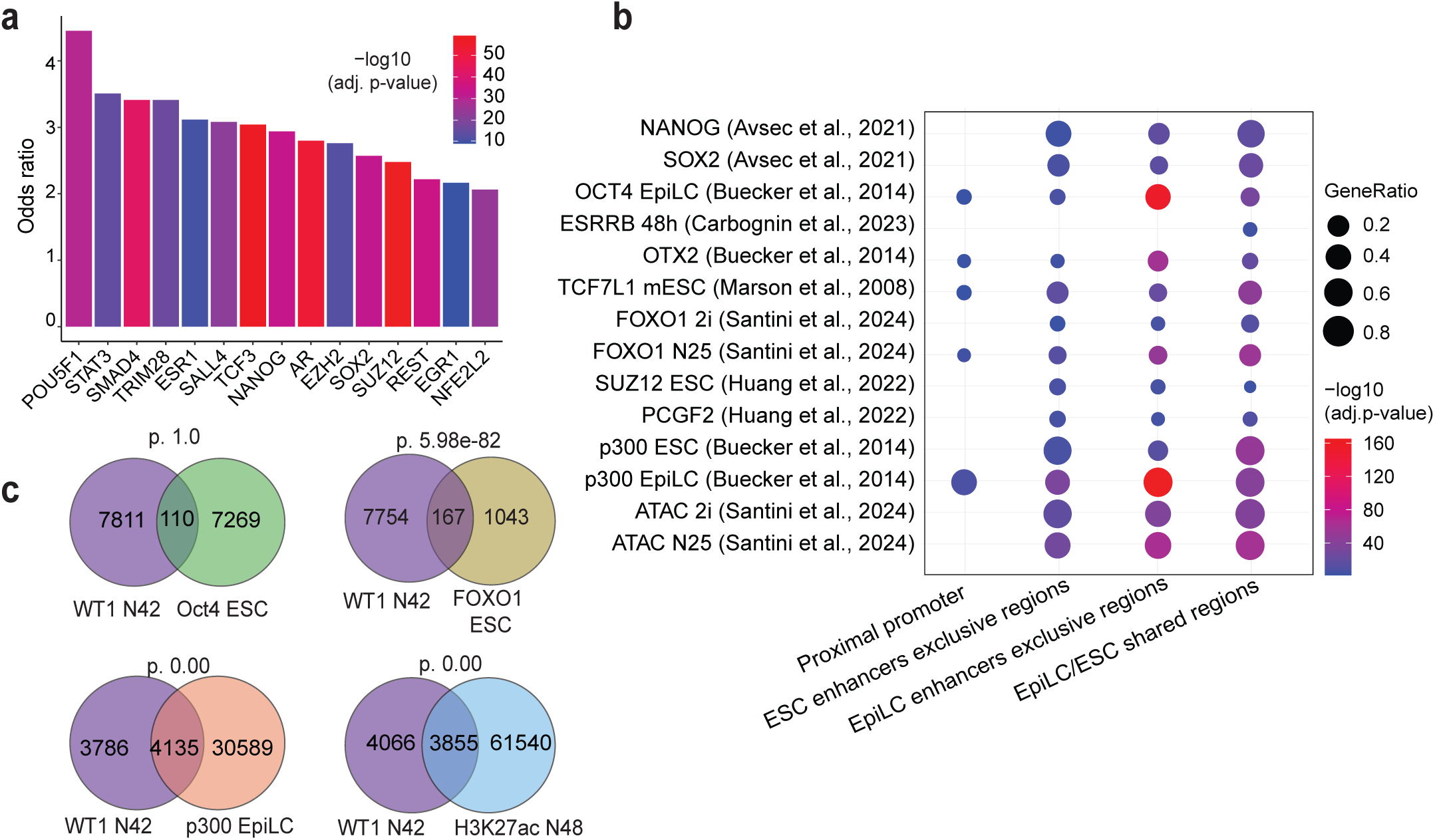
WT1 co-occupies active post-implantation regulatory elements and cooperatively enhances gene activation. a. Top 10 transcription factors whose ChIP-X–derived target gene sets significantly overlap with the WT1 target genes bound at N42. Enrichment was computed using Enrichr with the ENCODE and ChEA Consensus TFs from the ChIP-X gene set library. Bars indicate odds ratio and fill color represents -log_10_ (adjusted P value). b. Pairwise overlap between WT1 ChIP–seq peaks and multiple reference datasets (ChIP–seq and ATAC–seq) across four genomic feature categories, as in Figure 3a. Overlap significance was assessed using hypergeometric tests. Dot colour indicates enrichment significance, and dot size represents gene ratio. Non-significant comparisons (FDR ≥ 0.05) are not shown. c. Venn diagrams showing overlap between WT1 peaks at N42 and ChIP–seq peaks for OCT4 (ESC), OTX2, FOXO1 (ESC), p300 (EpiLC) and H3K27ac (EpiLC). Overlap significance was assessed using hypergeometric tests (background n = 155,266).

**Supplementary Figure 4.**
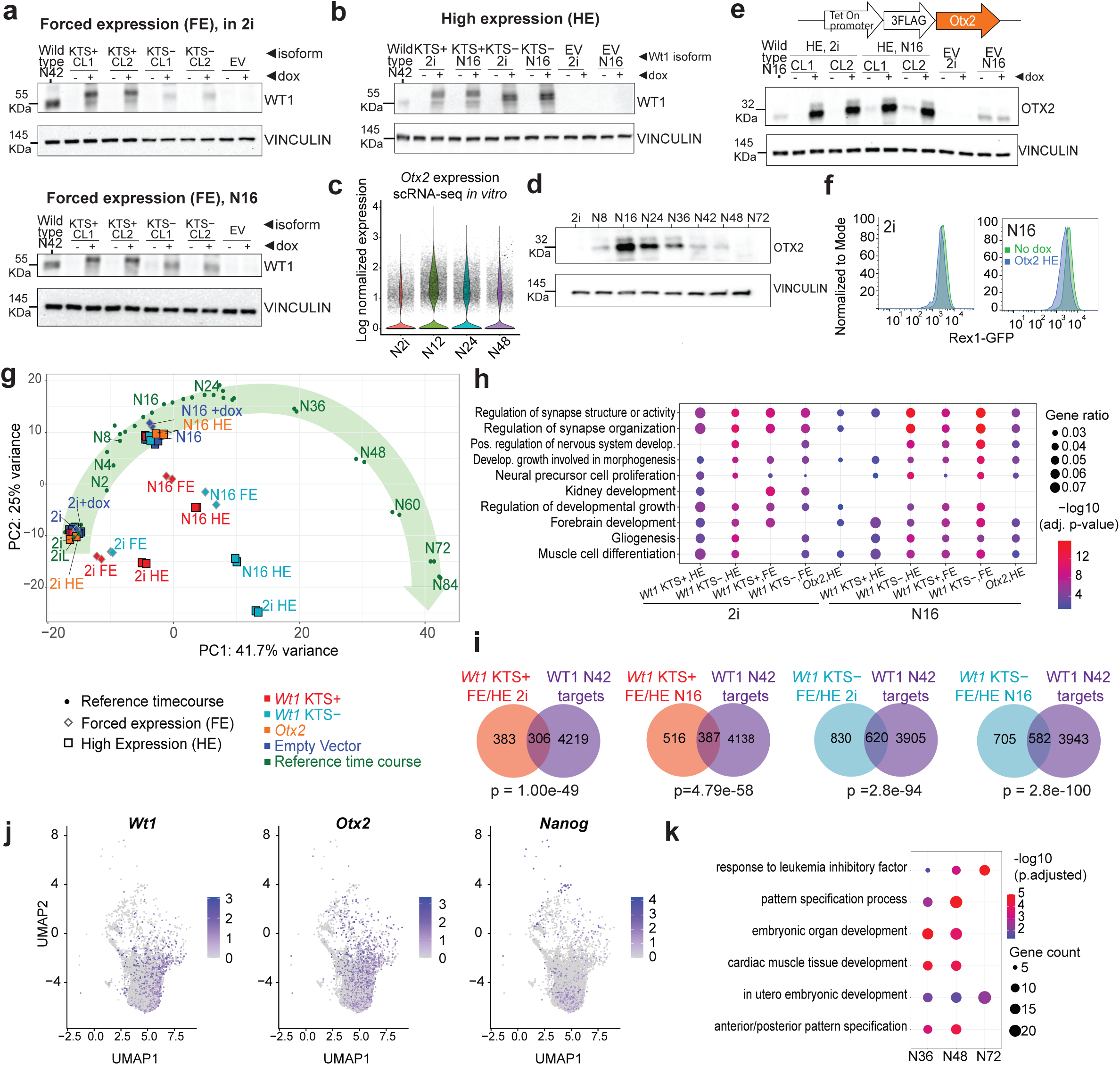
Forced early expression of *Wt1* leads to expedited acquisition of post-implantation identity a. *Wt1* forced expression (FE) conditions. Western blot of WT1 KTS+ an*d* WT1 KTS-in Rex1-GFP mESCs compared to endogenous WT1 peak expression at N42. CL1 and CL2 indicate independent clonal lines. b. Quantification of WT1 expression in high-expression (HE) conditions relative to endogenous N42 levels. c. *Otx2* expression across the in vitro single-cell RNA-seq time course, showing log-normalized expression during exit from naïve pluripotency d. OTX2 protein expression during exit from naïve pluripotency, showing peak expression at N16. e. Western blot defining OTX2 high-expression (HE) conditions following doxycycline titration. CL1 and CL2 indicate independent clonal lines. f. Representative Rex1-GFP flow cytometry profiles in 2i and N16 under *Otx2* high expression (clonal lines, n = 2). g. PCA of ComBat-corrected bulk RNA-seq integrating developmental time-course datasets^9,25^, with *Wt1* and *Otx2* forced expression samples. PC1 reflects developmental progression; *Wt1* forced expression shifts cells toward post-implantation states. Data points are biological replicates. h. Gene Ontology (Biological Process) enrichment of upregulated genes (padj ≤ 0.05, log_2_FC ≥ 0.5). Dot size indicates gene ratio; colour indicates -log_10_(FDR). i. Overlap between *Wt1*-induced transcriptional responses and WT1 ChIP-seq targets at N42. Venn diagrams show intersections for each isoform and condition (2i, N16). Enrichment significance was assessed using a hypergeometric test (background n = 21,859 protein-coding genes). j. Feature plots showing *Wt1*, *Otx2* and *Nanog* expression in the N48 single-cell RNA-seq dataset used for the analyses in Figure 4c. k. Dot plot showing representative significantly enriched Gene Ontology terms among genes downregulated in *Wt1* knockout cells during monolayer differentiation, using the DEG criteria defined in Fig. 4e.

**Supplementary Figure 5.**
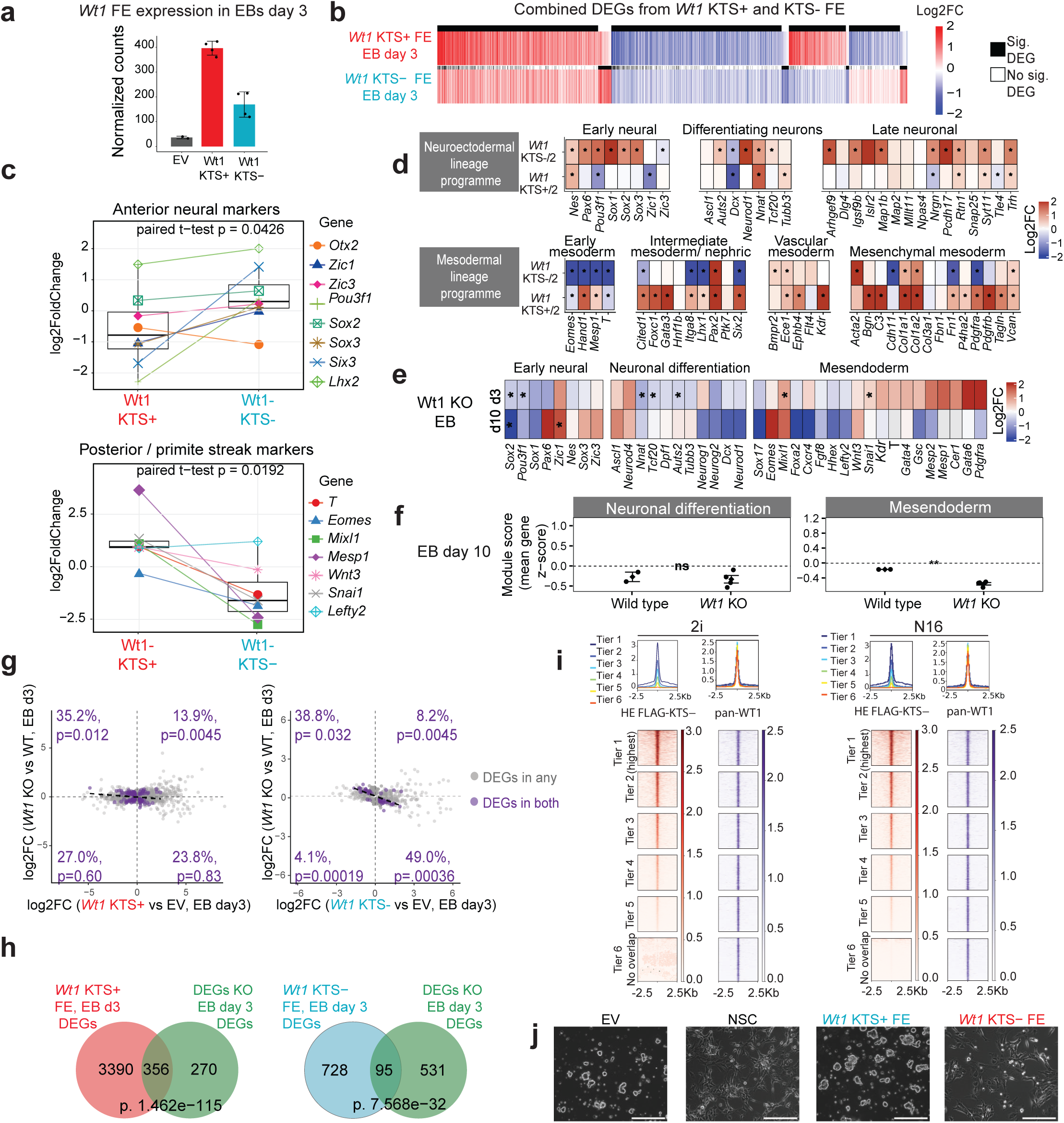
*Wt1* splice isoforms bias lineage programmes during early differentiation a. *Wt1* expression during embryoid body (EB) differentiation. mESCs carrying doxycycline-inducible *Wt1* KTS+ or *Wt1* KTS-transgenes were induced for three days, and *Wt1* transcript levels at EB day 3 were quantified from RNA-seq (DESeq2-normalized counts). b. Heatmap of log_2_ fold changes for the union of differentially expressed genes (adjusted p-value ≤ 0.1 in at least one comparison) following *Wt1* KTS+/1 or *Wt1* KTS-/1 forced expression (EB day 3, vs EV). Rows represent genes and columns represent conditions; colour scale indicates expression changes (-2 to +2). Side bars indicate statistical significance. c. Comparison of log_2_ fold changes for selected anterior neural and posterior primitive streak markers following *Wt1* isoform expression in EBs (day 3). P-values correspond to paired statistical tests. d. Lineage marker responses in second independent *Wt1* KTS+/2 and *Wt1* KTS-/2 clonal lines. Heatmaps show log_2_ fold changes at EB day 3 relative to control for the neuroectodermal and mesodermal lineage programme genes shown in Fig. 5c. Data are from two biological replicates per clonal line. Asterisks indicate adjusted p-value < 0.1. e. Heatmap showing transcriptional changes in *Wt1* knockout embryoid bodies at days 3 and 10. Log_2_ fold changes relative to wild type are shown for genes associated with early neural, neuronal differentiation and mesendoderm programmes. Genes are clustered separately within each programme based on their joint day 3 and day 10 expression profiles. f. Module-score analysis of formative pluripotency, neuronal differentiation and mesendoderm programmes in *Wt1* knockout embryoid bodies at day 10, calculated as described in Fig. 5d. Wild-type and *Wt1* KO samples are shown. g. Relationship between *Wt1* gain- and loss-of-function effects. Scatterplots show gene-wise log_2_ fold changes for *Wt1* isoform forced expression (x-axis) versus *Wt1* knockout (y-axis) for *Wt1* target genes. Genes significant in both contrasts are highlighted. Quadrant enrichment was assessed relative to an expected uniform distribution using χ² goodness-of-fit (χ² p = 0.0098 for *Wt1* KTS+ FE and χ² p = 2.11e-06 for *Wt1* KTS-FE) and binomial tests as indicated in the panels. h. Overlap between *Wt1* isoform forced-expression signatures and *Wt1*-KO DEGs in EBs (day 3). Venn diagrams show intersections between DEGs (adjusted p-value < 0.1) from gain and loss-of-function conditions. Enrichment significance was assessed using hypergeometric tests (background n = 21,859 protein-coding genes). i. FLAG ChIP-seq of *Wt1* KTS-under forced expression in naïve (2i) and differentiation-permissive (N16) conditions. Peaks shared with endogenous WT1 (N42) were stratified into binding tiers based on FLAG signal intensity; heatmaps show signal distribution across peak centres. j. Morphological comparison of neural stem cell (NSC) differentiation following *Wt1* isoform forced expression. Representative brightfield images of empty vector control, NSCs generated using the standard differentiation protocol, and cells subjected to *Wt1* KTS+ or *Wt1* KTS-forced expression followed by culture in NSC medium are shown. Scale bar, 200 μm.

**Supplementary Figure 6.**
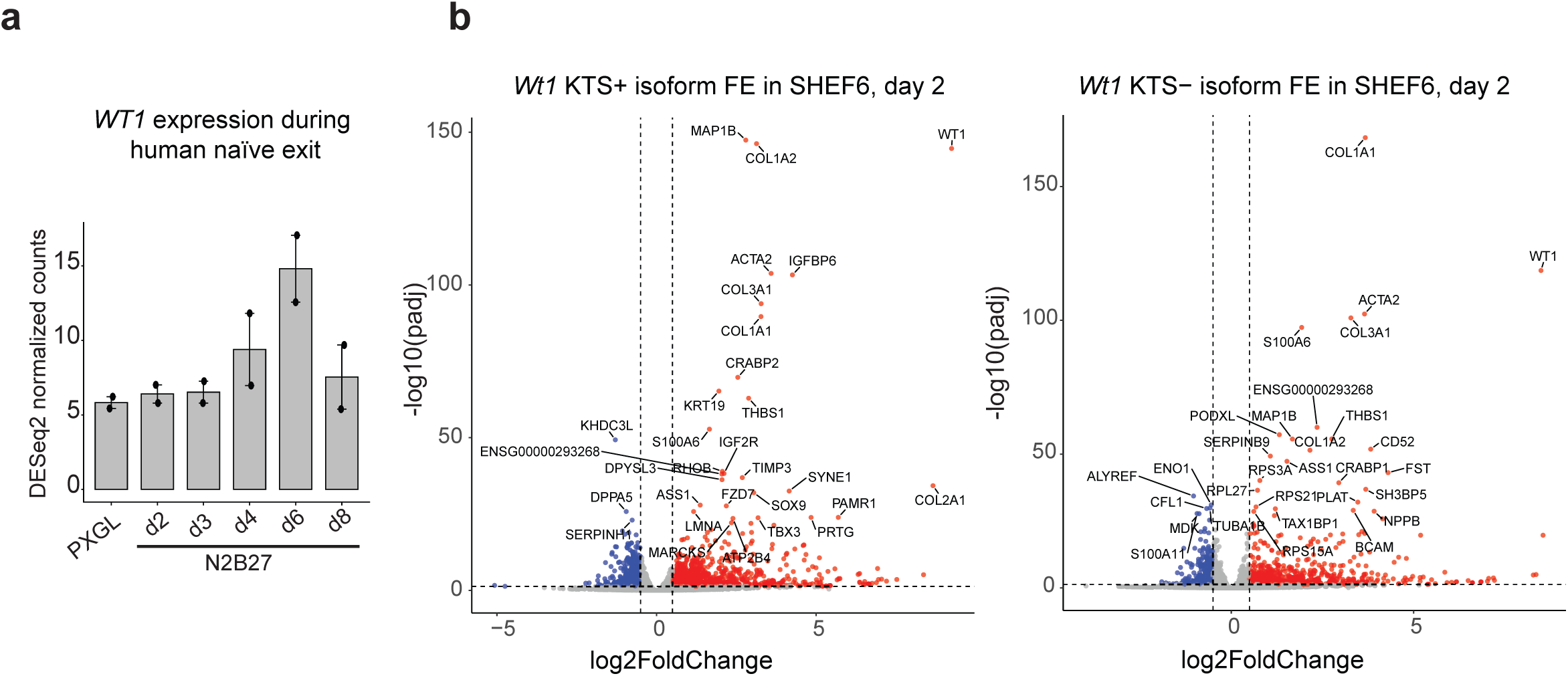
*Wt1* isoform programmes show cross-species transcriptional conservation. a. *WT1* expression during human naïve exit. WT1 expression in PXGL-Y naïve human ESCs and following transition to N2B27 for 2, 3, 4, 6 and 8 days, using publicly available RNA-seq data^41^. Bars represent mean DESeq2-normalized counts, black dots indicate individual biological replicates, and error bars indicate standard deviation. b. Volcano plots showing differential gene expression at day 2 of differentiation following *Wt1* KTS+ or *Wt1* KTS-forced expression in human SHEF6 cells relative to empty vector controls. Significantly upregulated and downregulated genes are highlighted, and selected top-ranked DEGs are labelled.

**Supplementary Figure 7.**
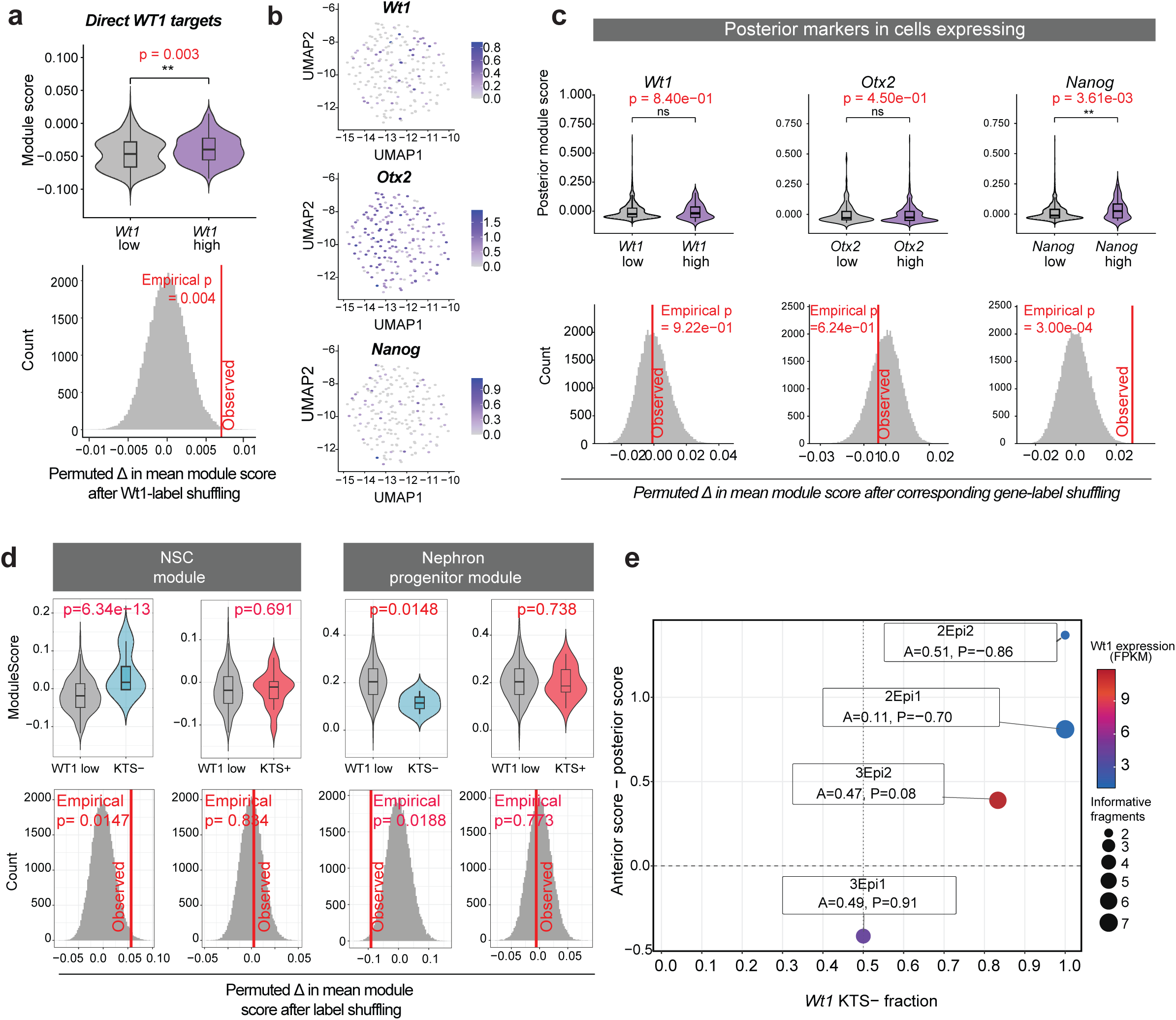
*Wt1* expression and splice composition align with lineage-biased transcriptional states in the E5.5 epiblast a. Violin plot showing per-cell module scores for a *Wt1*-associated transcriptional programme (595 genes upregulated upon *Wt1* induction and bound by WT1 at N42) in E5.5 epiblast cells from the integrated peri-implantation single-cell atlas. *Wt1*-high and *Wt1*-low cells are compared. Permutation histograms show null distributions generated by shuffling group labels (50,000 iterations); the vertical line indicates the observed difference in mean module score (Δ) between the two groups. b. Feature plots showing *Wt1*, *Otx2* and *Nanog* expression in E5.5 epiblast cells from the integrated peri-implantation single-cell RNA-seq atlas used in Figure 7. The plots illustrate the distribution of cells with detectable expression of each gene within the E5.5 epiblast population. c. Violin plots and permutation analyses comparing posterior module scores between *Wt1*-low and *Wt1*-high, *Otx2*-low and *Otx2*-high, and *Nanog*-low and *Nanog*-high cells. For each comparison, group labels were shuffled 50,000 times to generate null distributions; vertical lines indicate observed differences (Δ). d. Isoform-resolved analysis of *Wt1*-associated transcriptional bias. Violin plots and permutation tests compare cells with predominant KTS+ or KTS-representation with *Wt1*-low cells for anterior NSC and posterior nephron progenitor module scores using dataset from Mohammed et al.^26^. Null distributions were generated by label shuffling (50,000 iterations); vertical lines indicate observed Δ. e. Relationship between *Wt1* KTS splice-isoform balance and anterior-posterior transcriptional bias in epiblast spatial samples. The *Wt1* KTS-fraction was calculated from strict informative *Wt1* KTS splice-junction fragments, while anterior and posterior module scores were derived from gene-wise z-scored spatial expression values within each embryo. The y-axis shows the anterior-minus-posterior module score, point size indicates the total number of informative KTS fragments, and point colour indicates *Wt1* FPKM. Sample labels indicate the individual anterior (A) and posterior (P) module scores. The four informative E5.5 epiblast spatial samples were 2Epi2 (KTS+=0, KTS-=2), 2Epi1 (KTS+=0, KTS-=7), 3Epi2 (KTS+=1, KTS-=5) and 3Epi1 (KTS+=2, KTS-=2). Only epiblast samples containing informative KTS fragments are shown.

